# Genetic structure at national and regional scale in a long-distance dispersing pest organism, the bird cherry–oat aphid *Rhopalosiphum padi*

**DOI:** 10.1101/829986

**Authors:** Ramiro Morales-Hojas, Asier Gonzalez-Uriarte, Fernando Alvira Iraizoz, Todd Jenkins, Lynda Alderson, Tracey Kruger, Mike J. Hall, Alex Greenslade, Chris R. Shortall, James R. Bell

## Abstract

Genetic diversity is determinant for pest species’ success and vector competence. Understanding the ecological and evolutionary processes that determine the genetic diversity is fundamental to help identify the spatial scale at which pest populations are best managed. In the present study, we present the first comprehensive analysis of the genetic diversity and evolution of *Rhopalosiphum padi*, a major pest of cereals and a main vector of the barley yellow dwarf virus (BYDV), in Great Britain. We have used a genotype by sequencing approach to study whether i) there is any underlying population genetic structure in this long distant disperser pest at a national and regional scale; ii) the populations evolve as a response to environmental change and selective pressures, and; iii) the populations comprise anholocyclic lineages. Individual *R. padi* were collected using the Rothamsted Insect Survey’s suction-trap network at several sites across England between 2004 and 2016 as part of the RIS long-term nationwide surveillance. Results identified two genetic clusters in Great Britain that mostly paralleled a North – South division, although gene flow is ongoing between the two subpopulations. These different groups do not correspond to sexual and asexual types, sexual reproduction being predominant in Great Britain, and could correspond to ecotypes. Results also show that there is migration with gene flow across Great Britain, although there is a reduction between the northern and southern sites with the Southwestern population being the most genetically differentiated. There is no evidence for isolation-by-distance and other factors like primary host distribution could influence the migration patterns. Finally, results also show no evidence for the evolution of the *R. padi* population, and it is demographically stable despite the ongoing environmental change. These results are discussed in view of their relevance to pest management and the transmission of BYDV.

## Introduction

Insect pests are responsible for the loss of up to 20% of the major grain crops in the world with some models predicting that this will increase by 19 – 46% driven by a 2°C rise of the average global surface temperature (Deutsch et al., 2018). The efficacy and sustainability of new smarter control programmes will be contingent on the genetic variation of pest populations (Powell, 2018). This is because genetic variation determines the potential for adaptation and evolution of populations (i.e. how likely is that insecticide resistance will evolve; Hawkins, Bass, Dixon, & Neve, 2019), and virus transmission dynamics as different genotypes differ in their vector competence (Jacobson & Kennedy, 2013). Evolutionary and ecological factors such as gene flow and distribution of hosts affect genetic variation in populations, which influences the adaptive responses to environmental change and selective pressures (Caprio & Tabashnik, 1992; Davis & Shaw, 2001; Dong, Li, & Zhang, 2018).

Thus, the study of the genetic diversity across a species range can be used to infer ecological and evolutionary aspects of pest organisms that are difficult to study directly in wild populations, such as the levels of gene flow between populations and migration pathways, the population size and demographic responses to different environmental processes, or the ecological factors influencing the distribution of genetic diversity and adaptation of populations (Lushai & Loxdale, 2004; Roderick, 1996). For these reasons, the study of the geographic and spatial distribution of the genetic diversity is critical for developing preventive strategies to control and manage pest and pathogens. Therefore, this information will improve our capacity to monitor and forecast pest dynamics, which is fundamental for the development of new, more efficient smart crop protection approaches.

Aphids are major pests of crops and vectors of some of the world’s major plant viruses (Nault, 1997). They have a complex life-cycle, which includes asexual reproduction, host alternation and winged/wingless morphs (Van Emden & Harrington, 2017), and this variation has been suggested to play an important role in their success (Loxdale, Edwards, Tagu, & Vorburger, 2017). Many aphid species have a wide range of hosts, and species range from generalists like *Myzus persicae* to host specialist that can comprise host-specific strains with varying degrees of genetic divergence, like the case of *Acyrthoshipon pisum* (Blackman & Eastop, 2000; Peccoud, Maheo, De La Huerta, Laurence, & Simon, 2015; Peccoud, Ollivier, Plantegenest, & Simon, 2009). In addition, aphids show great plasticity in terms of reproductive type at the intraspecific level and lineages in some species range from sexual to obligate parthenogenetic (Blackman, 1974; Leather, 1993). This results in a great capacity to adapt to different environmental conditions; thus, it has been shown that populations of holocyclic species can remain reproducing parthenogenetically during winter in regions where temperatures are mild. As a result of the climatic response, there is a correlation between the proportion of asexuality and geography in some of the aphid pest species (Blackman, 1974; Llewellyn et al., 2003; Simon et al., 1999). This variation in life-cycle is central to their abundance and, importantly, this diversity also plays a central role in the transmission of viruses to crops. It is the winged aphids that transfer viruses between fields, however, at the field level it is the wingless individuals that spread the infection between plants (Jepson & Green, 1983; Ribbands, 1964). In holocyclic species, virus transmission in crops occur during the parthenogenetic phase, and the virus cycle is interrupted in winter by the return of the gynoparae (asexual females that produce the sexual females) to the primary host, usually a woody plant not suitable for the virus survival. In milder regions where individuals can survive reproducing parthenogenetically all year round, the virus transmission cycle in the crops will continue through the winter. Therefore, it is fundamental for virus risk evaluation to monitor the holocyclic and anholocyclic lineages. The variation in life cycle leaves a signature in the genome of populations, and the use of population genetic approaches can be used to infer the predominant mode of reproduction in populations (Halkett, Simon, & Balloux, 2005).

The bird cherry–oat aphid, *Rhopalosiphum padi*, is one of the major pests of cereals in the temperate regions, and it is a main vector of the barley yellow dwarf virus (BYDV) that causes cereal yield losses of between 20% to 80% (Vickerman & Wratten, 1979). *Rhopalosiphum padi* is a heteroecious species alternating between the primary host *Prunus padus* (bird cherry tree) and the secondary host, cereals and other grasses. While *R. padi* is predominantly holocyclic undergoing sexual reproduction in winter on the primary host, in some regions of continental Europe and perhaps even in southern Britain, mild winters favour permanent asexual reproduction on the secondary grass host. As a result, a cline in the proportion of parthenogenetic clones related to winter temperature has been described in France; in addition, sexual and asexual forms are genetically differentiated although some gene flow occurs through the occasional generation of males by asexual clones (Delmotte, Leterme, Gauthier, Rispe, & Simon, 2002; Martinez-Torres, Moya, Hebert, & Simon, 1997). In the UK it has also been suggested an increase of anholocyclic clones towards the south (Williams & Dixon, 2007). Despite its pest status across Europe, the genetic population structure and levels of gene flow have only been comprehensively studied in France. Here, sexual clones show high levels of gene flow across their range, indicating long distance migration, a so-called “long-distance disperser” (Delmotte et al., 2002). In Great Britain, *R. padi* is one of the most abundant species in the Rothamsted Insect Survey (RIS) 12.2 m suction traps, particularly in autumn during the migratory phase when individuals fly in large numbers to colonise the primary host, *P. padus*. Such migration suggests the “long-distance disperser” status, previously suggested in Britain given the homogeneous geographic genetic structure observed in alloenzyme studies (Loxdale & Brookes, 1988). The long-distance migration in the UK mostly occurs in the gynoparae individuals as a result of their migration to the sparsely dispersed primary host, *P. padus* (Tatchell, Parker, & Woiwod, 1983; Tatchell, Plumb, & Carter, 1988). Thus, the host plant distribution influences the migratory behaviour and ultimately the population genetic diversity of the species (Loxdale & Brookes, 1988).

In this study, we have analysed the distribution of the genetic variation in time and space of *R. padi* in Great Britain to provide further insight into key aspects of this pest ecology and evolution. We demonstrate that GBS approaches can detect population structure at national and regional scale in species that are long distant migrants that would show, *a priori*, a weak structure signal. Thus, we have analysed the population structure, gene flow levels and inferred the predominant mode of reproduction in the English populations. We have also analysed the proportion of sexual and asexual reproductive types in females collected during the autumn migration to confirm the genetic results. This study will be relevant to improve the management and control of cereal viruses vectored by this pest species, such as BYDV.

## Materials and Methods

### Samples

316 individuals of *R. padi* from the RIS biological archive collected in seven locations (Starcross, Wye, Writtle, Hereford, Preston, York and Newcastle; Figure 1) in 2004, 2007, 2010, 2013 and 2016 (Table 1 and Supplementary Table S1) were used in the present study. Samples in the biological archive of the RIS were preserved at room temperature in a solution containing 95% ethanol and 5% glycerol, although for this study the 2016 samples were stored at −20°C two weeks after collection rather than being committed to the archive.

**Figure 1.**
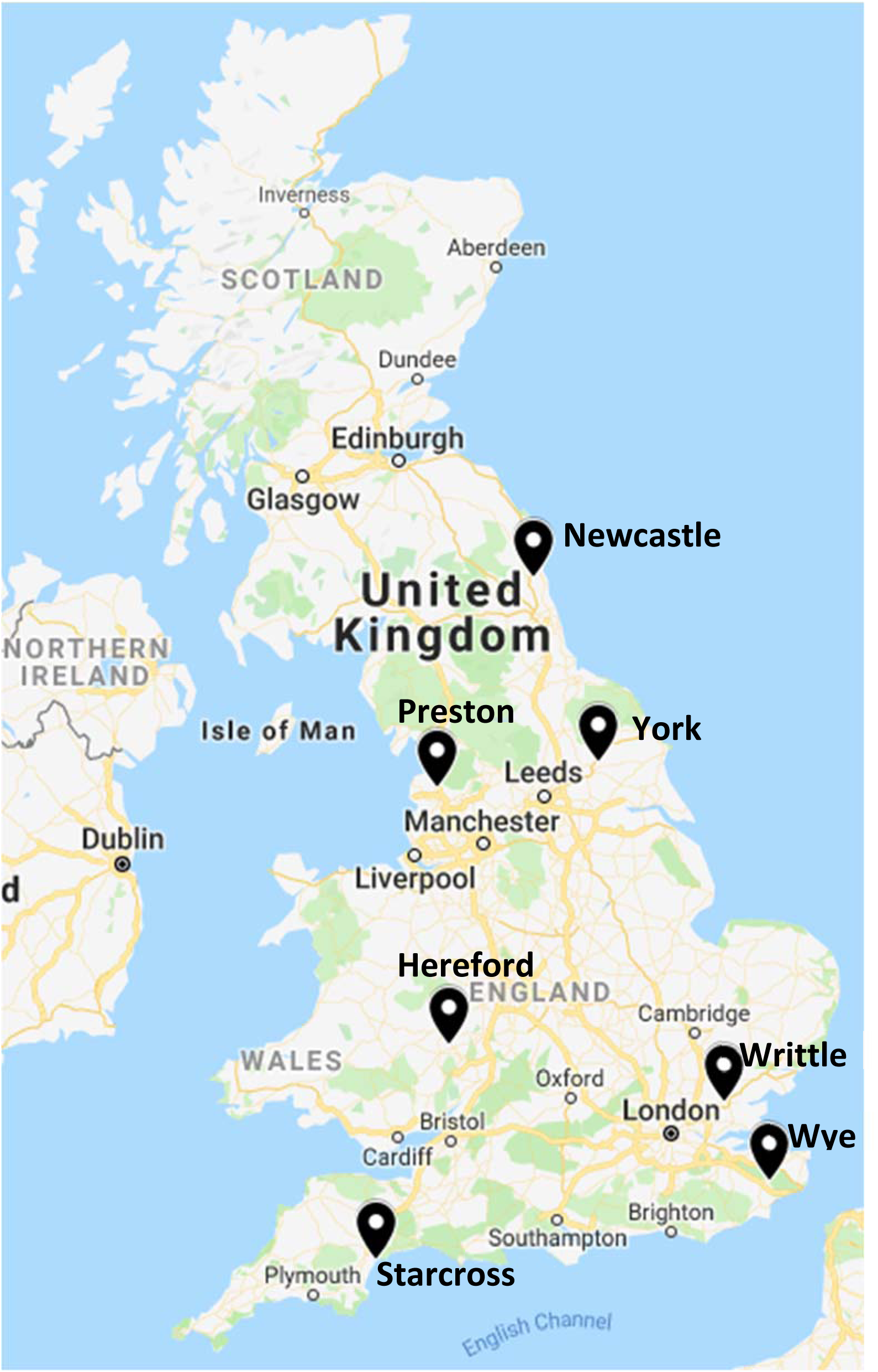
Map of Great Britain showing the location of the suction traps from which the samples were used.

**Table 1.** Summary of number of samples used for DNA extraction, kits used and the mean amount of DNA obtained per aphid.

|  | **2004** |  | **2007** |  | **2010** |  | **2013** |  | **2016** |  |
| --- | --- | --- | --- | --- | --- | --- | --- | --- | --- | --- |
| **Sites** | B&T | MK | B&T | MK | B&T | MK | B&T | MK | B&T | MK |
| **Starcross** | - | 10 | - | 10 | 20 | 10 | 20 | 10 | 26 | - |
| **Wye** | - | - | - | - | - | - | - | - | 15 | - |
| **Preston** | - | - | - | - | 10 | - | 14 | - | 16 | - |
| **Newcastle** | - | 10 | - | 10 | 10 | 10 | 10 | 10 | 15 | 5 |
| **Hereford** | - | - | - | - | - | - | - | - | - | 25 |
| **Writtle** | - | - | - | - | - | - | - | - | - | 23 |
| **York** | - | - | - | - | - | - | - | - | - | 27 |
| **Mean DNA yield (ng) (s. d.)** | - | 182.50 | - | 280.87 | 57.86 | 149.78 | 34.19 | 160.56 | 325.38 | 433.04 |
| **Standard Deviation** | - | 125.07 | - | 337.54 | 81.86 | 181.35 | 50.55 | 261.73 | 303.93 | 243.52 |

Note: B&T - QIAGEN's Blood & Tissue kit; MK - QIAGEN's DNA Micro kit.

### DNA extraction

DNA was extracted from single individuals using two commercial kits, QIAGEN’s DNeasy Blood and Tissue kit or QIAamp DNA Micro kit, to identify the method that provided better DNA yield and quality from the archive samples. DNA extractions were done following the manufacturer’s protocol but with the following modifications: individual aphid specimens were placed in 1.5 ml microcentrifuge tubes and submersed in liquid nitrogen prior to homogenisation with a sterile pestle; 180 µl of buffer ATL was added afterwards to the homogenised sample. Samples that were extracted with the Blood & Tissue kit were incubated with 5 µl of RNase A (100 mg/ml) for 30 minutes at 37 °C, followed by a 3 hours’ incubation with 20 µl of Proteinase K at 56 °C in a shaker at 180 r.p.m. The DNA Micro kit samples were instead incubated overnight at 56 °C with 20 µl of Proteinase K in a shaker at 180 r.p.m. but without a previous incubation with RNase A. Carrier RNA was added to Buffer AL as recommended in the DNA Micro kit manual for small samples, at a final concentration of 5 ng/µl of carrier RNA. The elution of the DNA was done with pre-warmed (56 °C) buffer AE, leaving the column stand for 10 minutes at room temperature before centrifugation; a second elution was performed using the first eluate. DNA yield was quantified using Qubit’s dsDNA assay.

### Genome sequencing and assembly

In order to improve the available genome of *R. padi*, long reads were generated using a MinION (ONT). DNA was extracted from a pool of four individuals of *R. padi* using QIAGEN’s DNeasy Blood and Tissue Kit. A total of 2.9 µg of DNA was used to prepare a genomic library following the ONT 1D ligation library protocol SQK-LSK108. A total of 1 µg of genomic library was loaded on a FLO-MIN107 flowcell and the sequencing run was performed with live base calling option on using MinKNOW v1.11.5.

Reads that passed the quality control were used in combination with Illumina reads publicly available (PRJEB24204) (Thorpe, Escudero-Martinez, Cock, Eves-van den Akker, & Bos, 2018a) to assemble the genome using MaSuRCA-3.2.8 (Zimin et al., 2013; Zimin et al., 2017). The resulting assembly was evaluated for completeness using Benchmarking Universal Single-Copy Orthologs (BUSCO v3.0.2) with the arthropoda orthoDB v9 set of genes, containing 1066 BUSCOs, and the insecta orthoDB v9 dataset of 1658 BUSCOs (Simao, Waterhouse, Ioannidis, Kriventseva, & Zdobnov, 2015). Gene prediction was done with Augustus v3.3.1 (Stanke, Diekhans, Baertsch, & Haussler, 2008) using the same trained set of genes used by Thorpe *et al*. (2018a) and RNAseq exon and intron hints. These hints were created by mapping RNAseq data available for *R. padi* PRJEB9912 and PRJEB24317 (ERR2238845 – ERR223884) (Thorpe, Cock, & Bos, 2016) to the genome assembly using the splice aware aligner STAR v2.4.0h (Dobin et al., 2013). Hints were created from the aligned bam and wig files using bam2hints (for the intron hints) and wig2hints (for the exon hints); these were merged to run Augustus. Gene models were annotated using Blast2GO 5.0.22 (Conesa et al., 2005). Blast was run using the arthropoda database.

### Genotyping of samples

A subsample of the aphid DNA samples was genotyped to investigate the population genetic diversity of *R. padi* in England. The samples sequenced were collected from Starcross (n = 73), Newcastle (n = 67), Preston (n = 35), Wye (n = 15), Writtle (n = 10) and York (n = 10) during March – October in 2004, 2007, 2010, 2013 and 2016 (Table 2; Supplementary Table S2). To increase the amount of DNA obtained from individual aphids, which in most cases was lower than 500 ng, whole genome amplification (WGA) was performed using Qiagen’s Repli-G UltraFast Mini Kit. The WGA reactions consisted of an equal volume of DNA (1 µl, 2 µl or 4 µl depending on the DNA template concentration) and buffer D1 (prepared following the Kit’s manual), which was incubated at room temperature for 3 minutes; after this incubation, an equal volume of buffer N1 was added (2 µl, 4 µl or 8 µl) to the reaction. This was mixed by flicking and spun down before adding 15 µl of reaction buffer and 1 µl of REPLI-g UltraFast DNA Polymerase. Reactions were incubated 2 hours at 30 °C followed by a 3 minutes incubation at 65 °C. DNA quantification was done using Qubit’s dsDNA BR assay. In some instances, a second WGA using 1 µl of whole genome amplified-DNA as template was performed when the DNA amount obtained after a first WGA was not enough for sequencing.

**Table 2.**
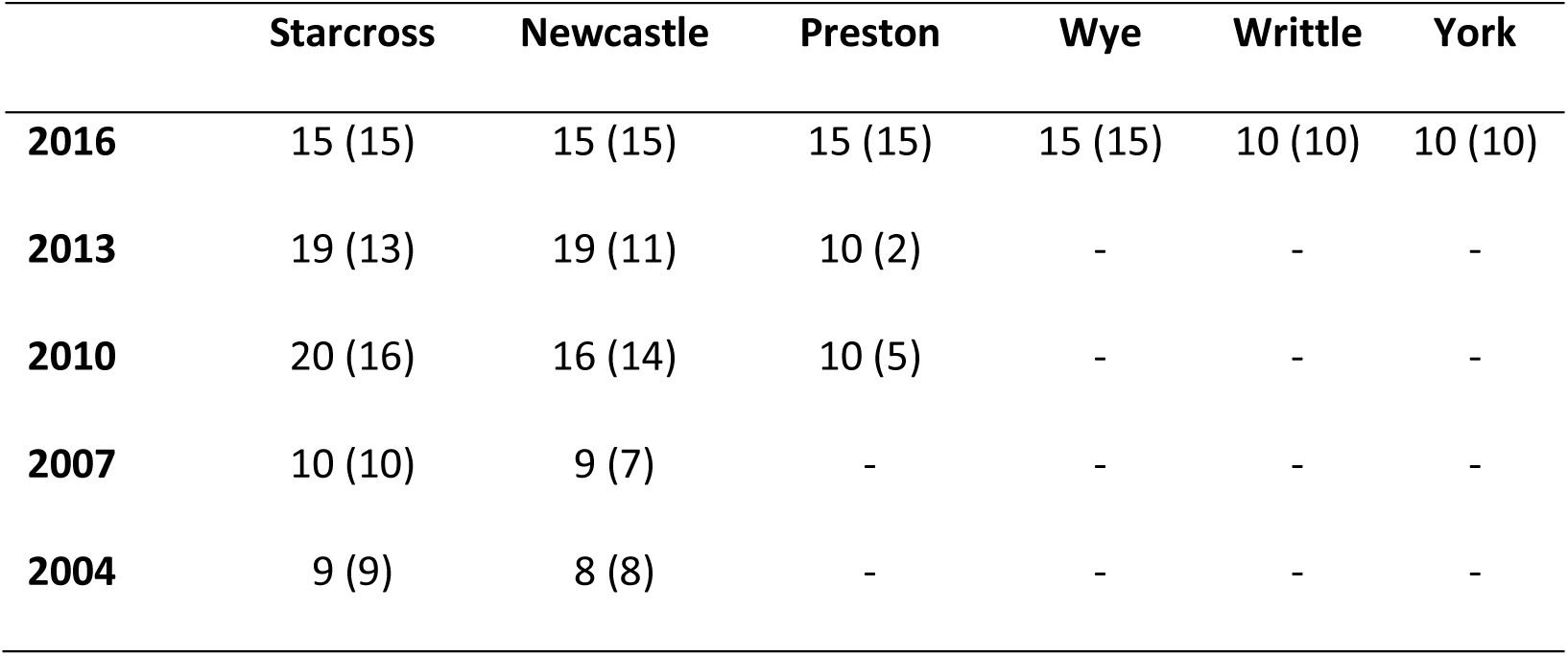
Number of samples genotyped using GBS per population and year; the number of successfully sequenced samples is shown in brackets.

|  | Starcross | Newcastle | Preston | Wye | Writtle | York |
| --- | --- | --- | --- | --- | --- | --- |
| **2016** | 15 (15) | 15 (15) | 15 (15) | 15 (15) | 10 (10) | 10 (10) |
| **2013** | 19 (13) | 19 (11) | 10 (2) | - | - | - |
| **2010** | 20 (16) | 16 (14) | 10 (5) | - | - | - |
| **2007** | 10 (10) | 9 (7) | - | - | - | - |
| **2004** | 9 (9) | 8 (8) | - | - | - | - |

Samples were genotyped using genome-wide single nucleotide polymorphisms (SNPs) identified following a genomic reduced representation sequencing method (genotype-by-sequencing, GBS). Sequencing of Writtle and York samples was done separately to those from the other four locations. At least 500 ng of genomic DNA per sample were digested with *Mse I* (primary digestion enzyme) and *Hae III* + *EcoRI* (secondary digestion enzymes), which were shown to provide an appropriate level of digestion and fragment size with an *in silico* evaluation of the *Acyrthosiphon pisum* genome. The library preparation and sequencing was performed following the standard Illumina pair-end (PE) protocol. PE sequencing of 150 bp was performed on an Illumina HiSeq platform. The GBS protocol was outsourced commercially.

Reads quality was assessed with FastQC v0.67 and mapped to the *R. padi* genome assembled in this study using BWA-MEM v0.7.16.0 (Li, 2013). Duplicates were removed using MarkDuplicates v2.7.1.1 and indels were realigned with BamLeftAlign v1.0.2.29-1. Variant calling was carried out with FreeBayes v1.0.2.29-3 (Garrison & Marth, 2012). The resulting SNPs from FreeBayes were annotated using snpEff v4.0. These tools were run using Galaxy v17.05 (Afgan et al., 2016).

### Analyses of population structure

SNPs called with FreeBayes were filtered using VCFtools v0.1.14 (Danecek et al., 2011) before the markers were used in subsequent analyses. Different filtering schemes were used to obtain a dataset that maximised the quality of the SNPs and genotypes while minimising the missing data at marker and individual levels (Supplementary Table S3), as recommended by O’Leary, Puritz, Willis, Hollenbeck, and Portnoy (2018).

The population structure was investigated using the Bayesian genetic clustering algorithm implemented in Structure 2.3.4 (Pritchard, Stephens, & Donnelly, 2000). We used the admixture model with correlated frequencies. To detect any potential subtle genetic structure, we ran Structure with the sampling locations set as priors (locprior = 1); this model has the power to detect a weak structure signal and does not bias the results towards detecting genetic structure when there is none. The *K* parameter was tested for values ranging from 1 to 6 with 10 simulations for each. We used 100,000 samples as burn-in and 200,000 samples per run for the Monte Carlo Markov Chain (MCMC) replicates. Parameter convergence was inspected visually. We ran the Structure simulations using a multi-core computer with the R package ParallelStructure (Besnier & Glover, 2013). The number of *K* groups that best fitted the dataset was estimated using the method of Evanno, Regnaut, and Goudet (2005) using Structure Harvester Web v0.6.94 (Earl & Vonholdt, 2012). Cluster assignment probabilities were estimated using the programme Clumpp (Jakobsson & Rosenberg, 2007) as implemented in the webserver CLUMPAK (Kopelman, Mayzel, Jakobsson, Rosenberg, & Mayrose, 2015). The genetic diversity measures of the populations were estimated using Arlequin 3.5.2.2 (Excoffier, Laval, & Schneider, 2005). Genetic variation among populations was investigated using an Analysis of the Molecular Variance (AMOVA) with 10000 permutations using Arlequin 3.5.2.2. We used hierarchical AMOVA to test the following population structures: i) differentiation between the North (Newcastle, Preston and York) and South (Wye, Starcross and Writtle) regions; ii) differentiation between genetic clusters identified with Structure; iii) differentiation of Newcastle populations from different years (2004, 2007, 2010, 2013 and 2016); iv) Starcross samples by year (2004, 2007, 2010, 2013 and 2016); and, v) differentiation of populations in different seasons (spring, summer and autumn) in the North (Newcastle and Preston), South (Wye and Starcross), and all locations together. Population pairwise divergence was investigated using F_ST_, and the significance was evaluated with 10000 permutations in Arlequin. We ran a Mantel test as performed in Arlequin to test for correlation between the genetic distances (F_ST_) and the geographic distance between sampling locations estimated using Google maps. Demographic events such as population expansion or bottleneck were inferred using Tajima’s D (Tajima, 1989) and Fu’s F_S_ (Fu, 1997) as estimated in Arlequin.

Phylogenetic trees of haplotypes were constructed using Maximum Likelihood (ML) with RAxML 8.2.12 (Stamatakis, 2014) run in the server CIPRES (Miller, Pfeiffer, & Schwartz, 2010). RAxML was run with 1000 bootstrap inferences with subsequent ML search using the gtrgamma model. The Lewis correction for ascertainment bias was implemented as it is the appropriate model for binary datasets that include only variable sites (as it is the case of SNPs) (Leache, Banbury, Felsenstein, de Oca, & Stamatakis, 2015; Lewis, 2001).

### Identification of reproductive types in autumn migrating females

The identification of virginoparae (asexual females producing asexual progeny) and gynoparae (asexual females that produce sexual females) individuals migrating in autumn is an indirect method to estimate the proportion of parthenogenetic and sexual reproducing lineages. This is because gynoparae females migrate to the primary host to produce the sexual forms while the virginoparae are migrating to winter cereals and other grasses to reproduce parthenogenetically throughout winter. The RIS has been monitoring the autumn migration of *R. padi* (mid-September to mid-November) and recording the number of virginoparae and gynoparae since 1995. For this, aphids are collected alive using the Rothamsted suction trap and the method of Lowles (1995) is used to differentiate the reproductive form of migrating *R. padi* females. This method relies on the colour differences of embryos when females are dissected in ethanol, with embryos from gynoparae females being green-yellow and those from virginoparae red-brown. In the present study, we have estimated proportion of sexual and asexual autumn migrating females in 2004, 2007, 2010, 2013 and 2016 for consistency purposes with the genetic analyses.

## Results

### Genome sequencing, assembly and annotation

The genome of *R. padi* has been recently sequenced (Thorpe et al., 2018a). To improve the continuity of the published assembly, genomic DNA from *R. padi* was sequenced using a MinION (Oxford Nanopore Technology, ONT). The number of reads obtained that passed the quality control was 1,111,646, with a distribution of lengths ranging from 63 bp to 74,319 bp (median length 3006 bp, mean length 3625; Figure S1). These long reads were used together with the short Illumina reads generated by Thorpe et al. (2018a) to assemble a new genome using MaSuRCA. The resulting assembly has a length of 321,589,008 bp, which is similar to the length of the previous genome version (Table 3). However, the continuity of the new assembly is much improved (2,172 versus 15,616 scaffolds) and the N50 has been increased from 116,185 bp in the Thorpe at al. (2018a) genome to 652,723 in the new assembly (Table 3). The completeness of the new version of the genome is higher, having identified 96.8% and 93.9% complete BUSCO genes of the Arthropoda and Insecta dataset, respectively (Table 3). The number of gene models identified in the new genome assembly is 26,535, which is similar to the number of genes in the previous version; 77% of these genes had a BLAST hit in the Arthropoda database (Table 3). 90% of the top-hit species were from other aphid species, and the similarity of the BLAST hits ranged from 33-100% (Supplementary Figures S2 and S3).

**Table 3.** Genome assembly statistics of the Illumina assembly of Thorpe et al. (2018) and the short and long-reads hybrid assembly obtained with Masurca.

|  | Illumina genome | Masurca genome |
| --- | --- | --- |
| **Assembly size (Mb)** | 319 | 321 |
| **Scaffolds** | 15,616 | 2,172 |
| **Scaffold N50 (bp)** | 116,185 | 652,723 |
| **Longest scaffold (bp)** | 616,405 | 4,088,110 |
| **N (bp)** | 54,488 | 0 |
| **GC (%)** | 27.8 | 27.8 |
| **BUSCO (complete, duplicated, fragmented, missing)** | 82%, 8.1%, 7.8%, 9.4% | Insecta: 93.9%, 3.6%, 1.4%, 4.7%<br>Arthropoda: 96.8%, 3.1%, 0.7%, 2.5% |
| **Genes** | 26, 286 | 26,535 |
| **BlastP hit** | 20,368 (77%) | 20,481 (77%) |

### Genotype by sequencing of historical samples

DNA yield was low in general (Table 1, Supplementary Table S1), but there was a significant difference in the amount of genomic DNA obtained between samples collected in different years (Supplementary Results, Supplementary Figure S4). We performed whole genome amplification of the DNA for most samples to obtain the required DNA amount for GBS (Supplementary Table S2). In the case of 21 samples, it was necessary to perform a second WGA reaction using as template 1 µl of the DNA product from the first WGA reaction. Of the 210 samples chosen for genotyping, 175 provided enough DNA for library construction and sequencing (Table 2). Of the 175 samples, 80 were from 2016, 26 from 2013, 35 from 2010, 17 from 2007 and 17 from 2004 (Table 2). The average number of reads obtained per sample was 3,708,552, with an average mapping to the *R. padi* genome of 65.67% (Supplementary Table S4). The number of reads obtained and mapped are significantly higher for the samples from 2016 (Mann-Whitney test *P* < 0.05 after Bonferroni correction for all pairwise comparisons), while there are no significant differences in the sequencing results from older samples. The total number of SNPs identified with FreeBayes when all samples were analysed together was 2,287,871. The proportions of missing data per individual and per locus for the complete dataset ranged from 0 to 1, reflecting the high variability in the quality of the sequencing results (Supplementary Figures S5 and S6). Different filtering schemes were applied to the SNP dataset to identify the one that maximised the quality of the called SNPs and resulted in the highest number of called genotypes in the maximum number of individuals (Supplementary Table S3).

### Genetic diversity of R. padi in England

When all the samples were included in the analyses, filtering scheme 7 (FS7) resulted in 4,802 SNPs in 86 individuals (Table 4). The final proportion of missing data per locus was < 5% and per individual was < 45% (Figure 2), hence the subsequent analyses were carried out using the FS7 dataset.

**Table 4.** Sequential steps done in the filtering schemes (FS) that provided the best dataset to be used in subsequent population analyses. The order of rows indicates the sequential filters applied to the data.

| **FILTER** | **All Samples**<br>**FS7** | **Samples Newcastle**<br>**FS2** | **Samples Starcross**<br>**FS2** |
| --- | --- | --- | --- |
| **MISSING DATA** | max-missing > 50%<br>remove-indels | remove-indels<br>max-missing > 50% | remove-indels<br>max-missing > 25% |
| **LOW-CONFIDENCE SNP CALL** | mac > 3<br>minQ > 20<br>minDP > 3 | min-meanDP > 5<br>mac > 3<br>minQ > 20 | min-meanDP > 5<br>mac > 3<br>minQ > 20 |
| **MISSING DATA** | imiss < 60%<br>max-missing > 90% | imiss < 95%<br>max-missing > 80% | imiss < 95%<br>max-missing > 70% |
| **LOW-CONFIDENCE SNP CALL** | minDP > 5 |  |  |
| **MISSING DATA** | max-missing > 95% |  |  |
| **ADDITIONAL FILTERS** | thin 2000 | thin 2000 | thin 2000 |
| **SNPs** | 4802 | 3186 | 907 |
| **INDIVIDUALS** | 86 | 36 | 31 |

*Note:* minDP – include only genotypes greater or equal to the value; min-meanDP – retain loci with a minimum mean depth of the given value; minQ – include sites with quality above the value; mac – include sites with minor allele count greater or equal to the value; max-missing – retain loci that have been successfully genotyped in the given proportion of individuals; imiss – retain individuals with a proportion of missing data smaller than the value; thin – remove loci that are closer than the given number of bp.

**Figure 2.**
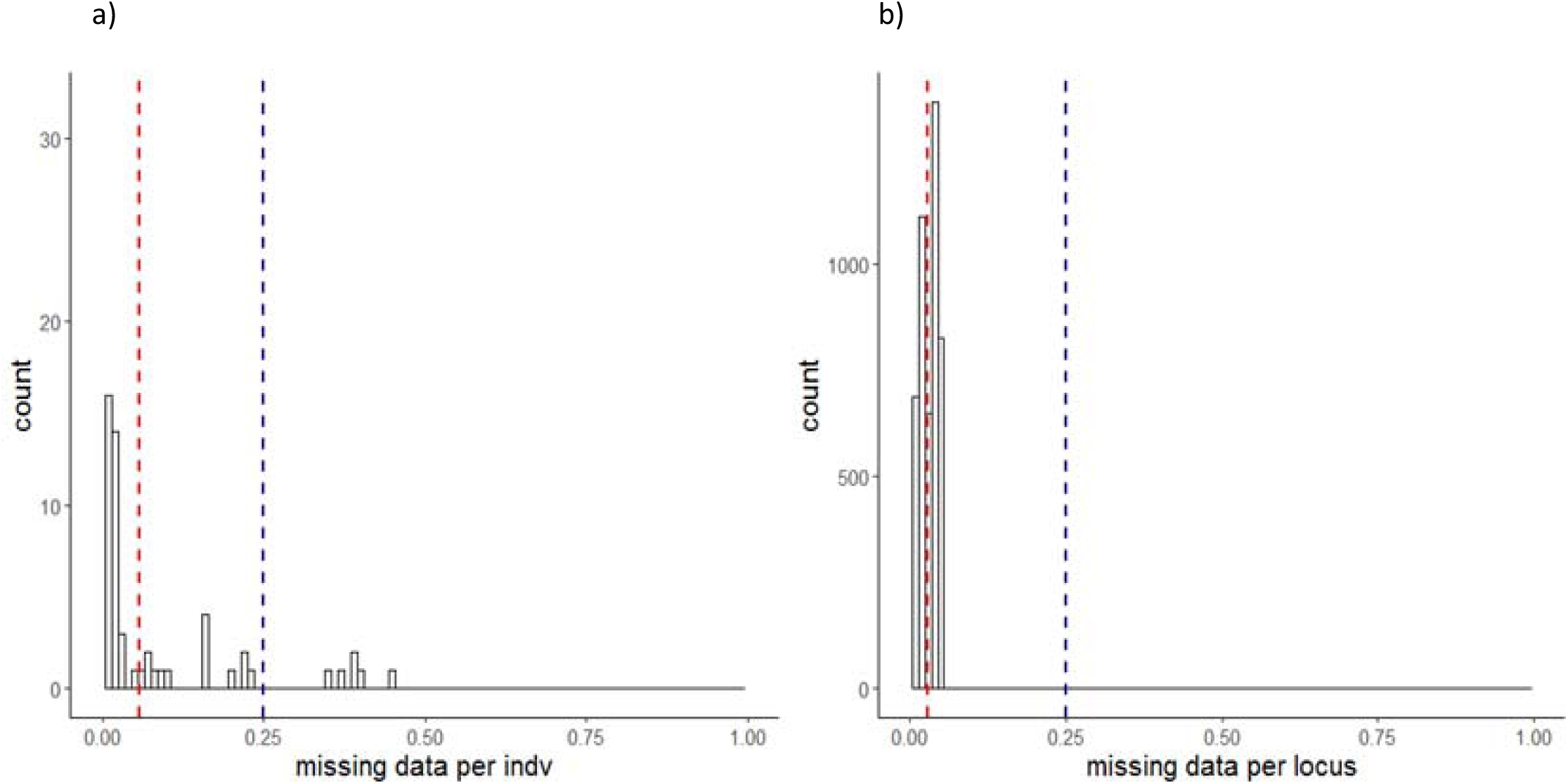
Distribution of the missing data per individual (a) and locus (b) in all samples after applying the filter scheme 7 (FS7). Vertical dashed lines in red correspond to the mean missing data.

A total of 172 different haplotypes were observed in the population. The nucleotide diversity in all samples was π 0.21 and θ_w_ 0.17 (Table 5). The gene diversity (also called expected heterozygosity, He) was higher than the observed heterozygosity (He = 0.24, Ho = 0.13), and the inbreeding coefficient (F_IS_) was 0.42 (*P* = 0) (Table 5). 74% of the individual loci had significantly different He and Ho (*P* < 0.05), however, there was no significant deviation from the Hardy-Weinberg equilibrium (HWE) at the haplotype level when the SNPs were phased. The nucleotide diversity (π and θ_w_) was similar across all populations except for a higher π in Wye and a lower θ_w_ in York (Table 5). The observed heterozygosity (Ho) averaged over all SNPs was higher in the southern Starcross (0.20) and Wye (0.26) sites than in the northern populations at Newcastle (0.14), York (0.11) and Preston (0.16) (Table 5), but the lowest Ho was observed in the samples collected in Writtle (0.08) also a southern site. The lower Ho observed in York and Writtle could be the result of library effects resulting from having been genotyped separately from the other samples, which can bias the resulting datasets (Bonin et al., 2004; O’Leary et al., 2018). Nevertheless, PCA analysis does not show any clustering of the samples according to genotyping experiment, suggesting that there is no bias in the loci as a result of library effects (Supplementary Figure S7). The mean Ho was lower than the mean expected heterozygosity (He) in all locations and the inbreeding coefficient (F_IS_) observed in all sampling locations was positive and significant (Table 5). Nevertheless, there was no deviation from the HWE at haplotype level in any of the populations (*P* = 1), although 37% (Newcastle), 25% (Preston), 17% (Starcross), 13% (Wye), 30% (Writtle) and 21% (York) of individual loci deviated significantly from the HWE. The F_IS_ levels were higher in York and Writtle, which could be because the samples from these two locations were collected during a small period of two to three weeks in late July to early August (Supplementary Table S4) with an increased probability of sampling individuals from the same asexually reproducing clones, and thus biasing the result to a higher inbreeding coefficient.

**Table 5.** Genetic diversity estimates for the six sampling locations; all the samples; North and South populations; and, the genetic clusters (GC) identified by Structure analyses.

|  | **N** | **H** | **$\pi$** | **$\theta_w$** | **Ho** | **He** | **F<sub>IS</sub>** | **D** | **F<sub>S</sub>** |
| --- | --- | --- | --- | --- | --- | --- | --- | --- | --- |
| **Starcross** | 24 | 24 | 0.20 | 0.20 | 0.20 | 0.34 | *0.31* | 0.004 | 1.1 |
| **Wye** | 30 | 30 | 0.25 | 0.20 | 0.26 | 0.30 | *0.14* | 0.796 | 0.831 |
| **Writtle** | 20 | 20 | 0.21 | 0.18 | 0.08 | 0.34 | *0.74* | 0.738 | 1.610 |
| **Newcastle** | 54 | 54 | 0.19 | 0.20 | 0.14 | 0.25 | *0.38* | -0.062 | -1.237 |
| **Preston** | 24 | 24 | 0.19 | 0.21 | 0.16 | 0.29 | *0.43* | -0.266 | 1.076 |
| **York** | 20 | 20 | 0.18 | 0.15 | 0.11 | 0.33 | *0.67* | 0.656 | 1.409 |
| **All** | 172 | 172 | 0.21 | 0.17 | 0.13 | 0.24 | *0.42* | 0.80 | -13.48 |
| **South** | 74 | 74 | 0.22 | 0.18 | 0.17 | 0.27 | *0.36* | 0.82 | -2.32 |
| **North** | 98 | 98 | 0.19 | 0.19 | 0.12 | 0.23 | *0.45* | 0.15 | -4.85 |
| **GC North** | 120 | 120 | 0.19 | 0.17 | 0.12 | 0.23 | *0.44* | 0.49 | -7.44 |
| **GC South** | 52 | 52 | 0.21 | 0.17 | 0.23 | 0.31 | *0.20* | 0.81 | -0.97 |

*Note:* N – number of gene copies (2 x number of individuals), H – number of haplotypes, $\pi$ – nucleotide diversity (average proportion of pairwise differences over all loci), $\theta_w$ – nucleotide diversity (average of segregating sites across all loci), Ho – mean observed heterozygosity over all loci, He – mean expected heterozygosity (gene diversity) over all loci, and F<sub>IS</sub> – population specific inbreeding coefficient (significant values at 5% are shown in italics), D correspond to Tajima's D and F<sub>S</sub> to Fu's F<sub>S</sub> tests (F<sub>S</sub> significance is set to be $\leq 2\%$ level as recommended in the original paper).

Nevertheless, the levels of gene diversity (He) in Writtle and York were similar to those of the other populations, suggesting that a comparable range of genetic diversity was sampled at each site and that not just a restricted number of clones were migrating at the time of sampling. In the case of the other locations, the genotyped samples were collected from March to October (Supplementary Table S4), reducing the probability of re-sampling individuals from the same clones because the sampling period was spread over several months.

### *Geographic structure of* R. padi

Results from the Structure analyses with the FS7 dataset identified a *K* = 2 as the most probable number of genetic groups in the *R. padi* population in England (Supplementary Figure S8). These two genetic clusters corresponded broadly to the south and north regions (Figure 3), although there were individuals in these two regions that had a higher probability of genetic membership to the alternative genetic cluster. This admixture is more common in the samples of southern origin (Starcross, Wye and Writtle), in which more individuals had a higher probability of membership to the northern genetic cluster. The genetic differentiation (F_ST_) within *R. padi* inferred by a non-hierarchical AMOVA was low but significant (0.041, P = 0.007). The north – south genetic differentiation was further explored using analyses of the molecular variance (AMOVA) (Table 6A). This hierarchical AMOVA showed that 3.74% of the genetic variation could be attributed to geographic origin (north and south regions), although the F_CT_ (0.037) was not significant (*P* = 0.1); 3.22% of the genetic variation arose between populations within geographic clusters and 93.05% occurred within populations. The hierarchical AMOVA when individuals were grouped according to the genetic clusters identified in the Structure analysis showed that 17.20% (*P* = 0.001) of the genetic variation occurred between the two clusters, while the majority (79.24%, *P* = 0) still occurred within locations.

**Figure 3.**
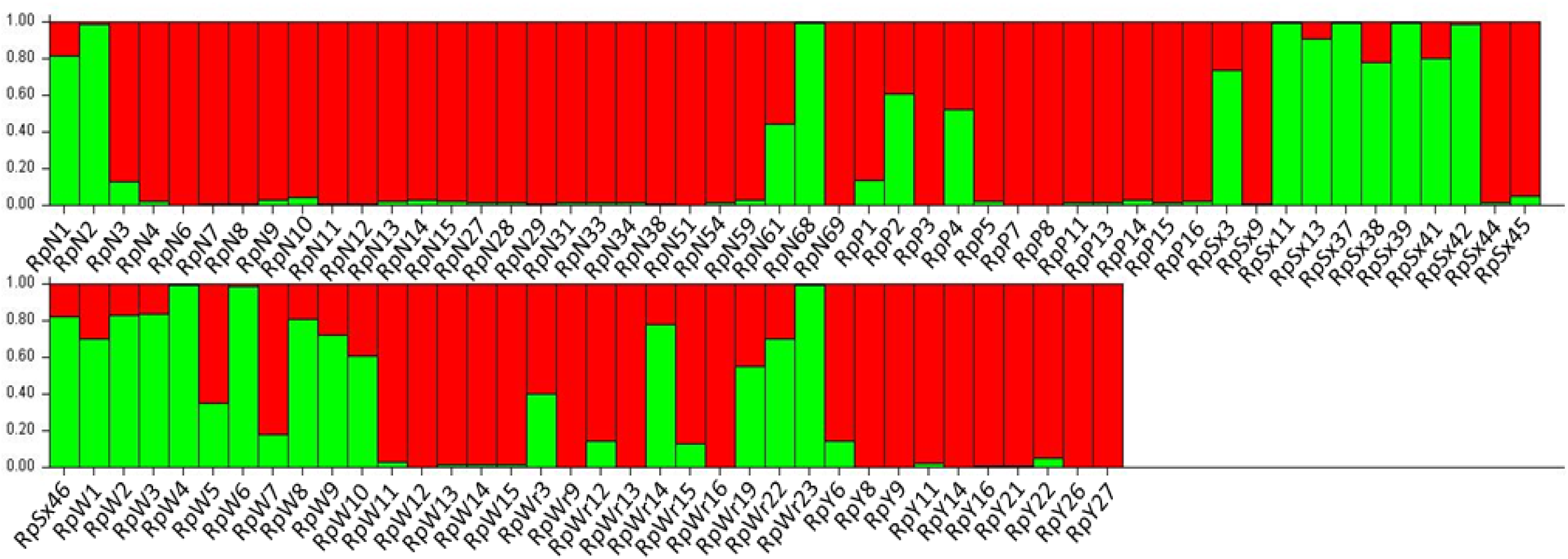
Bar plot resulting from Structure analysis when *K*=2 and sorted by population. The bars represent individuals and the colour of the bars represent the probability of membership to a certain population.

**Table 6.** Hierarchical AMOVA results for genetic structure of *R. padi*. A) two geographic clusters comprising individuals from the North (Newcastle, Preston and York) and the South (Starcross, Wye and Writtle); B) two genetic clusters as determined by the Structure analyses.

| Source of variation | d.f. | Sum of squares | Variance components | % variation | Fixation indices | P value |
| --- | --- | --- | --- | --- | --- | --- |
| **A** |  |  |  |  |  |  |
| Among North and South | 1 | 2704.904 | 19.832 Va | 3.74 | $F_{CT} = 0.0373$ | 0.1 |
| Among locations within North and South | 4 | 3802.365 | 17.077 Vb | 3.22 | $F_{SC} = 0.0334$ | 0 |
| Within locations | 166 | 82101.888 | 494.041 Vb | 93.05 | $F_{ST} = 0.0695$ | 0 |
| **B** |  |  |  |  |  |  |
| Among genetic clusters | 1 | 8037.878 | 99.332 Va | 17.20 | $F_{CT} = 0.1719$ | 0.001 |
| Among locations within clusters | 9 | 6788.132 | 20.589 Vb | 3.56 | $F_{SC} = 0.0430$ | 0 |
| Within locations | 161 | 73692.147 | 457.715 Vc | 79.24 | $F_{ST} = 0.2076$ | 0 |

The genetic differentiation levels, estimated as population pairwise F_ST_, were low to moderate and significant between the different sampled locations (Table 7). Starcross showed the highest differentiation with respect to the northern populations compared to others in the South (Wye and Writtle). There was no genetic differentiation within each region (Preston v. Newcastle F_ST_ = 0.01, *P* = 0.05; Wye v. Starcross F_ST_ = 0.005, *P* = 0.22) except for York in the north and Writtle in the south. Mantel test did not identify a significant relationship between genetic and geographic distance between sampling locations, suggesting that the differentiation is not the effect of isolation by distance. The genetic differentiation between the north and south regions was F_ST_ = 0.05 (*P* = 0); when the differentiation was estimated for the genetic clusters identified by Structure, the F_ST_ was higher (F_ST_ = 0.18, *P* = 0).

**Table 7.** Genetic differentiation between the sampled populations. Population pairwise F_ST_ values are shown below the diagonal. Significant F_ST_ are shown in italics.

|  | Newcastle | Preston | York | Starcross | Wye | Writtle |
| --- | --- | --- | --- | --- | --- | --- |
| Newcastle | - |  |  |  |  |  |
| Preston | 0.010 | - |  |  |  |  |
| York | *0.058* | *0.059* | - |  |  |  |
| Starcross | *0.081* | *0.096* | *0.166* | - |  |  |
| Wye | *0.037* | *0.042* | *0.095* | 0.005 | - |  |
| Writtle | *0.049* | *0.062* | *0.074* | *0.057* | *0.033* | - |

The estimated phylogenetic tree had little bootstrap support and the internal branches were in general short (Figure 4). Nevertheless, there is a weak clustering pattern that corresponded to the two genetic clusters identified by the Structure analysis. The southern group was paraphyletic with respect to the northern genetic clade, which is monophyletic and shows higher bootstrap support (64%). This weak support for the phylogenetic clades is expected because, despite the high level of genetic differentiation between the two genetic groups there is still gene flow between the locations. Estimates of the Tajima’s D and Fu’s F_S_ for the different populations did not deviate significantly from neutrality (Table 5), which is an indication of a stable demographic history. When these statistics were estimated for the complete dataset, Tajima’s D was not significant (D = 0.803, *P* = 0.75) while the value of Fu’s F_S_ was negative and significant (F_S_ = −13.479, *P* = 0.01), indicative of a significant departure from the neutral expectation. Similarly, when these tests were run for the two identified genetic clusters, F_S_ was negative and significant only for the genetic cluster comprising most individuals of a northern origin (F_S_ = −7.44, *P* = 0.02) whilst Tajima’s D was not significant. A negative value of F_S_ evidences an excess of alleles, which would be expected after a recent population expansion. Fu’s F_S_ is considered to be more sensitive to population expansion than the Tajima’s D statistic and is the best performing statistic for large sample sizes, but its sensitivity to recombination may result in significant values even if no expansion has taken place (Ramirez-Soriano, Ramos-Onsins, Rozas, Calafell, & Navarro, 2008). Thus, given the genome-wide nature of our dataset, it is likely that recombination is biasing the Fu’s F_S_ results.

**Figure 4.**
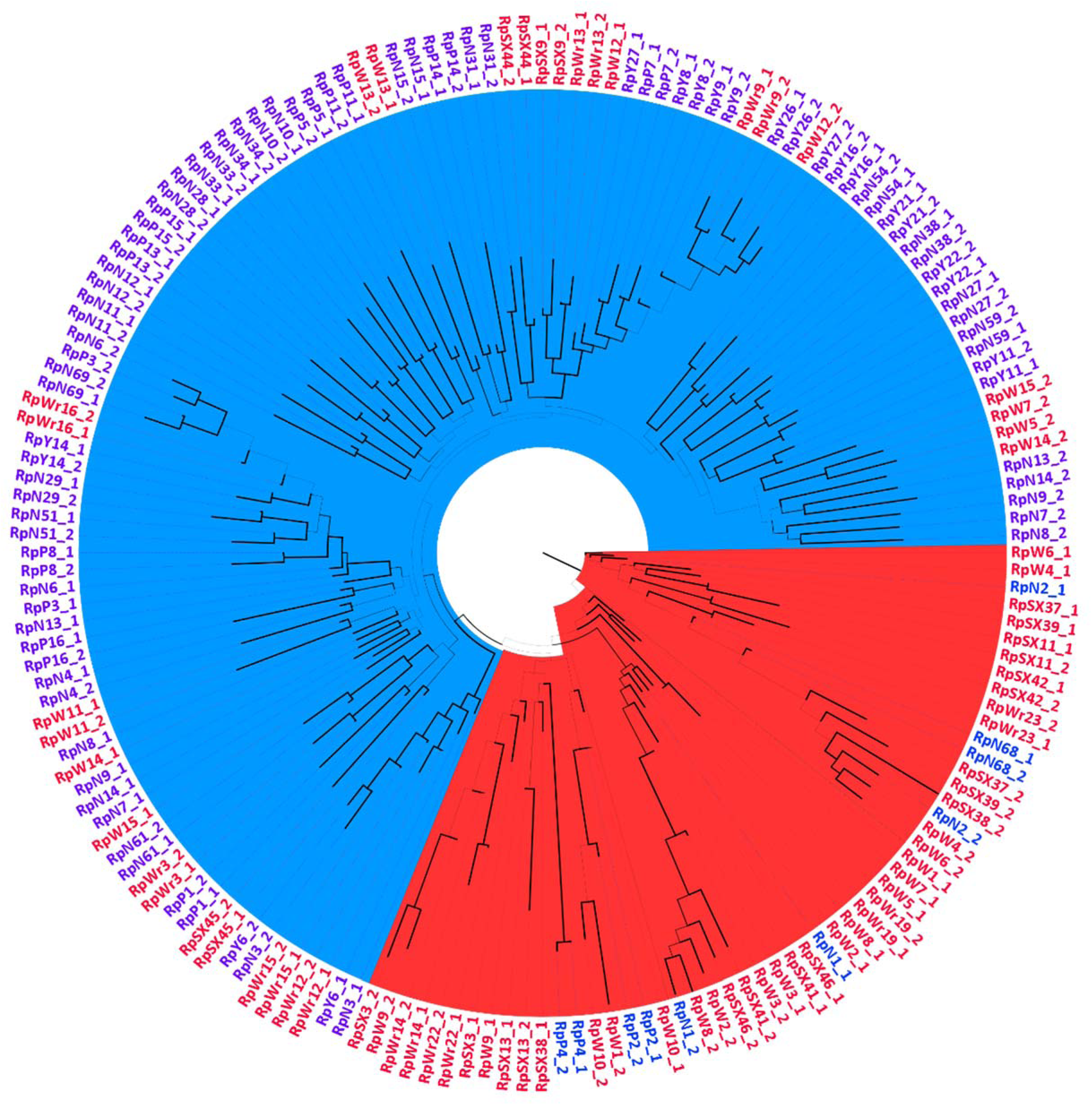
Midpoint rooted ML phylogenetic tree of haplotypes from the FS7 dataset. Branch line weight relates to bootstrap support, wider line corresponds to a bootstrap > 90%. Clades representing the genetic clusters identified with Structure, are highlighted with different colours (blue – northern cluster, red – southern cluster). Haplotypes from individuals collected in the southern sites are in red, those from the North are in blue.

### *Temporal differentiation of* R. *padi populations*

To explore the temporal differentiation of populations, we analysed separately the samples from two locations, Newcastle and Starcross, collected in 2004, 2007, 2010, 2013 and 2016. Different filtering schemes were used to identify the SNP dataset that maximised the quality and number of markers and individuals and minimised the missing data (Supplementary Table S3B and C). The best results were obtained with FS2 (Table 4), although the proportion of missing data per individual was still high in a few samples (Supplementary Figure S9). Thus, 7 and 13 individuals had more than 70% of the loci missing in the Newcastle and Starcross datasets, respectively; still, 29 individuals from Newcastle and 17 from Starcross had less than 50% of missing data. In the case of Starcross, no samples from 2007 were kept after filtering. In both populations, the pairwise differentiation (F_ST_) between years was low and non-significant (Table 8), indicating a lack of genetic differentiation within populations through time.

**Table 8.** Pairwise F_ST_ values between samples from Newcastle (A) and Starcross (B) collected in different years. Significant values are shown in italics. N shows the number of individuals in each year in the FS2 dataset.

**A)**
|  | N | 2016 | 2013 | 2010 | 2007 | 2004 |
| --- | --- | --- | --- | --- | --- | --- |
| 2016 | 15 | - |  |  |  |  |
| 2013 | 2 | -0.481 | - |  |  |  |
| 2010 | 8 | -0.037 | -0.284 | - |  |  |
| 2007 | 5 | 0.008 | -0.228 | 0.069 | - |  |
| 2004 | 6 | -0.201 | 0.084 | 0.0005 | 0.005 | - |

**B)**
|  | N | 2016 | 2013 | 2010 | 2007 | 2004 |
| --- | --- | --- | --- | --- | --- | --- |
| 2016 | 14 | - |  |  |  |  |
| 2013 | 5 | -0.095 | - |  |  |  |
| 2010 | 7 | -1.301 | -2.098 | - |  |  |
| 2004 | 5 | -0.727 | -0.457 | -0.602 |  | - |

Samples used in the present analyses were collected at different points of the season, resulting in genotyping of aphids sampled during the first flight to crops in April-May, during the mid-season in July-August and when aphids fly back to winter host in October. This allows the analysis of the genetic consequences in populations as the seasonal selective pressures change. Analyses have been carried out separately for all samples from the North and South using the FS7 dataset; York and Writtle were not included in these seasonal analyses because the samples were all collected in July. The genetic differentiation between spring, summer and autumn samples was significant for all pairwise comparisons except in the case of spring and summer in the South (Table 9). When the differentiation was estimated between location-season, the pattern becomes more complex (Supplementary Table S5). The autumn populations were significantly differentiated from spring and summer populations, except in Starcross. The spring and summer populations were not significantly differentiated in Preston and Wye, but have significant F_ST_ values in Newcastle (0.057, *P* = 0.01) and Starcross (0.079, *P* = 0.049). However, it should be noted that the seasonal analyses at population level could be biased by the different number of samples, which in some cases was low. It is interesting to note that the gene diversity varied throughout the season in all locations, although the variation pattern was different for each of the locations (Figure 5). In general, the genetic diversity as measured by the He was higher in the spring than in summer and autumn when the samples were grouped by geographic region (north and south). Nevertheless, the He in the south decreased only in the autumn sample, while in the north the He decayed from spring to summer and increased again in the autumn to a similar level to that of the southern population (Figure 5).

**Figure 5.**
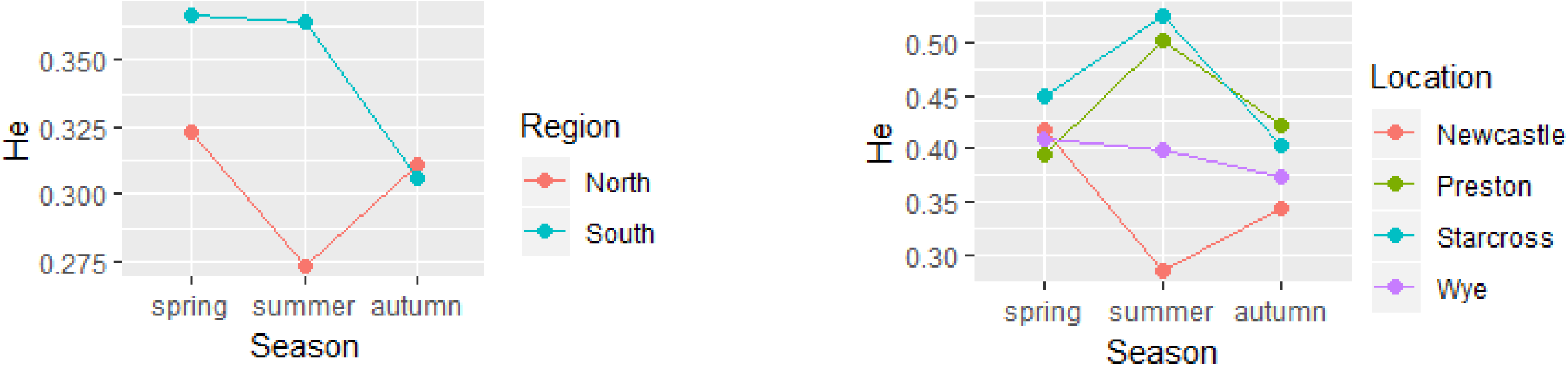
Plot of the genetic diversity calculated for the different regions and populations at different times of the season.

**Table 9.** Genetic differentiation (F_ST_) between samples collected in spring, summer and autumn in the North (A) and South (B) locations. Values in italics are significant. N shows the number of samples.

**A)**
|  | N | Spring | Summer | Autumn |
| --- | --- | --- | --- | --- |
| **Spring** | 9 | - |  |  |
| **Summer** | 18 | *0.026* | - |  |
| **Autumn** | 12 | *0.042* | *0.036* | - |

**B)**
|  | N | Spring | Summer | Autumn |
| --- | --- | --- | --- | --- |
| **Spring** | 9 | - |  |  |
| **Summer** | 8 | -0.005 | - |  |
| **Autumn** | 10 | *0.080* | *0.033* | - |

### Proportion of asexual lineages in R. padi population

The number and proportion of virginoparae and gynoparae females migrating in autumn collected from the 16 of September to the 14 of November in 2004, 2007, 2010, 2013 and 2016 are shown in Table 10. The proportion of virginoparae highly varied from year to year, with 2007 and 2013 being higher than 50%, while in 2010 it was approximately 2%. The average proportion of virginoparae across the analysed years is approximately 16%.

**Table 10.** Number and proportion of migrating females collected in the Rothamsted suction trap between the 16 of September and the 14 of November in 2004, 2007, 2010, 2013 and 2016 that were virginoparae and gynoparae.

| Year | 2004 | 2007 | 2010 | 2013 | 2016 | All |
| --- | --- | --- | --- | --- | --- | --- |
| **Virginoparae** | 35 | 96 | 10 | 13 | 29 | 183 |
| **Gynoparae** | 312 | 82 | 398 | 125 | 20 | 937 |
| **Total** | 347 | 178 | 408 | 138 | 49 | 1120 |
| **% Virginoparae** | 10.1 | 53.9 | 2.4 | 9.4 | 59.2 | 16.3 |

## Discussion

The present study represents the first longitudinal analysis of the population genetic diversity of *R. padi* using a genomic approach with samples from the biological archive of the RIS that go back to 2004. This study shows that reduced genome sequencing approaches can detect population genetic structure in highly migratory species like *R. padi* (Delmotte et al., 2002; Loxdale & Brookes, 1988), which *a priori* should show a weak signature of genetic differentiation, and help identify predominant mode of reproduction in the population, which is essential to understand plant virus transmission. This approach has also been able to detect weak signatures of seasonal genetic variation, which could be due to selection pressure changes.

The genome of *R. padi* was recently made publicly available (Thorpe et al., 2018a), which facilitates the identification of genome-wide markers. In the present study we have used long reads to improve the assembly and quality of the available genome. Thus, the number of scaffolds has been reduced more than 7x while maintaining a similar genome size as that of the previous version. In addition, the quality has increased as measured by the proportion of single ortholog genes identified, which has increased from 82% complete genes to 94% in the present assembly. Nevertheless, the number of genes annotated in the present genome is similar to that the previous version. The availability of a quality genome assembly has helped with the identification and analyses of genome-wide molecular markers for the population genetics analyses of this species in England.

The study has identified two genetic clusters that correspond broadly to a geographic differentiation between the south and the north locations. These two genetic clusters explain 17% of the genetic variation, which is higher than the source of variation between sexual and asexual populations in France that was shown to be 11% (Delmotte et al., 2002). The two clusters identified in England could also correspond to sexual and asexual forms; however, the inbreeding coefficient F_IS_ is positive and significant in both clusters indicating a deficiency in the observed heterozygosity, which is normally explained by factors like inbreeding or admixture of subpopulations (Wahlund effect). Instead, asexual reproduction results in an excess of heterozygotes and therefore a negative F_IS_, as observed in the French asexual population of *R. padi* (Halkett et al., 2005). In addition, the presence of individuals with mixed southern and northern genotypes are most likely the result of sexual reproduction. This is also apparent in the phylogenetic tree, which shows short internal branches with low bootstrap support. This uncertainty in the phylogenetic relationships is possibly due to incongruent phylogenetic signal across the SNP dataset, which could arise as a result of recombination (Brito & Edwards, 2009). Overall, the results of the present study suggest that the two genetic clusters identified in *R. padi* do not correspond to different reproductive forms. This finding is relevant in that the population of *R. padi* in England could comprise two different ecotypes, which could play different roles in the transmission of BYDV.

Genetic variation results indicate that sexual reproduction is dominant in the English populations of *R. padi*. This has been confirmed by the examination of the reproductive type of migrating females in autumn, which detects a low mean number of asexual lineages when averaged over all years analysed. However, there are years when the proportion of asexual females migrating in autumn is high. These could be either facultative parthenogens and therefore the asexual genetic signal would be lost when they reproduce sexually the following year. Alternatively, if they are obligate parthenogenetic lineages, the lack of genetic signal in the population would result from a high mortality in winter (Williams, 1987), maintaining the number of asexually reproducing lineages low in the population. This is relevant to BYDV transmission because it is during the asexual phase that *R. padi* becomes highly relevant as a vector of the virus, and asexual individuals can maintain the circulation of the virus in the winter cereal crops where they overwinter. As *R. padi* in Great Britain primarily reproduces sexually in the primary host, individuals are unlikely to maintain BYDV circulating during winter, reducing the possibility of outbreaks early in the following growing season. This situation, however, needs to be monitored as the proportion of asexually reproducing females varies from year to year, but also because the expected winter temperature rise due to climate change can reverse the situation and make anholocyclic lineages dominant in Great Britain.

Sexual populations of *R. padi*, *S. avenae* and other aphids have been shown previously to have a significant homozygosity excess (Delmotte et al., 2002; Hebert, Finston, & Foottit, 1991; Simon et al., 1999; Simon & Hebert, 1995). Four hypotheses have been proposed to explain this in *R. padi* (Delmotte et al., 2002): the presence of null alleles, which is unlikely the case in our dataset as filters have been applied to minimise the number of loci with missing genotypes; selection due to differential winter survival between lineages; inbreeding; and, allochronic isolation due to different timing in the production of sexual aphids between different lineages. Differential survival between lineages was discarded by Delmotte et al. (2002) due to their sampling scheme. Here, we can also rule out this explanation as natural selection would result in allele frequency variation and genetic differentiation between years, which is not observed in this study. Inbreeding linked to a patchy distribution of host plants was proposed to be the cause of an excess of homozygosity in *Melaphis rhois* (Hebert et al., 1991). In the case of *R. padi*, the distribution of the primary host *P. padus* is also patchy across Great Britain and specifically sparse in the south. The inbreeding coefficient is higher in the north, where individuals do not have to migrate long distances to find a *P. padus* tree to reproduce and overwinter, increasing the probability of related individuals mating. On the other hand, the individuals from the south will need to migrate longer distances as *P. padus* is rarer, decreasing the probability of inbreeding. However, the lack of significant genetic differentiation within the geographic regions indicate that the distribution of *P. padus* is not the only factor influencing the significant excess of homozygosity. While the distribution of the primary host is likely to influence the genetic variation of *R. padi*, further studies of the biology and diversity should be performed to determine the factors resulting in the observed pattern of deviation of HWE.

In addition to the two genetic clusters, it was observed that the levels of genetic differentiation varied geographically. There was no significant differentiation between the two northern-most locations (Preston and Newcastle) or the two southern-most (Wye and Starcross); but, there was a significant differentiation between the northern and southern sites with the Southwest being the most genetically differentiated population. The two locations of York and Writtle showed high genetic differentiation with respect to all the other populations, which was not expected given their geographic location. There was no correlation between the geographic and genetic distances between the locations, so there is no isolation by distance. This suggests that there are other factors than just geographic distance that are reducing the migration of individuals between the North and the South, and mostly in the Southwest. Ecological factors such as the distribution of the primary host, *P. padus*, is likely to influence the dispersal capacity of individuals although, as discussed above, it is probably not the only feature influencing the migration of this species. It should be noted that the two genetic clusters account for most of the differentiation within the species, and these correspond to the north and south regions. Thus, even when there is migration between the geographic regions as evidenced by individuals from each genetic cluster being collected in the alternative geographic region, the genetic differentiation is due to a reduction in gene flow between the two genetic forms. Landscape genetic analyses should be carried out with more dense sampling to determine the resistance surfaces that reduce migration between regions.

Climate change is shifting the distribution and reducing the genetic diversity in many species (Balint et al., 2011). In aphids, it has been observed that their phenology has changed during the last 50 years, with an earlier first flight (Bell et al., 2015). Nevertheless, there has been no significant change in their long-term populations size despite yearly cycles in abundance. The availability of historical *R. padi* samples from the last 20 years in the Rothamsted Insect Survey has allowed the study of the evolution of its populations through time. The analyses have found no significant genetic differentiation between 2004 and 2016 populations, indicating no evolution of the population in Great Britain; moreover, there is no signature of demographic bottleneck or expansion either. Thus, there is no indication that climate change has affected the abundance of this pest in Britain, which is consistent with the observations of Bell *et al*. (2015); there is also no evidence for a reduction in the genetic diversity of the populations through time, which could suggest that they are stable and resilient to changing climatic conditions.

Another aspect of temporal dynamics is the seasonal change in populations. This study has identified low but significant differentiation levels between samples collected in spring, summer and autumn, with this latter being the most differentiated. There is also a variation in the observed heterozygosity and genetic diversity from spring to autumn. Heterozygosity has been observed to decrease from the primary to the secondary host populations of *R. padi* in Canada (Simon & Hebert, 1995), and this phenomenon was explained by either the presence of admixture of homozygous migratory individuals and resident clones or by a selective disadvantage of heterozygotes. In both cases, the genetic diversity (He) of populations does not necessarily decrease. Thus, when there is an admixture of heterogeneous populations (migrants and resident clones) in the secondary host, different alleles would come together in the same population; in the case of clonal selection against heterozygotes, the expected heterozygosity is not necessarily reduced as homozygotes for different alleles could remain in the population. However, in this study, the He also decreases from the primary to the secondary, so there must be alternative explanations that would result in the reduction of both the observed heterozygosity and the genetic diversity in the crop. The strong selection that adaptation to a new host plant and insecticide use impose on populations could explain such pattern. It has been observed in *Myzus cerasi* that the transfer of individuals from the primary (cherry trees) to the secondary host (cleavers) in laboratory conditions resulted in a survival rate of 10-20% (Thorpe, Escudero-Martinez, Eves-van den Akker, & Bos, 2018b). The two hosts of *R. padi* are similar to those of *M. cerasi* in that the primary host is a tree, the bird cherry, and the secondary host is herbaceous, cereals; therefore, it is possible to expect a low survival rate in the species when changing hosts. This would explain the genetic differentiation between the spring, summer and autumn migrating samples as well as the reduction in genetic diversity (He) in relation to the spring samples. In addition, insecticide use on crops during the growing season can also result in a further reduction of the heterozygosity and the allelic diversity during summer. Nevertheless, no insecticide resistance has been observed in populations of *R. padi* in the UK and it would be expected a larger reduction in genetic diversity in the autumn migrating population than that observed in the study. The possibility remains that a large proportion of the *R. padi* population remains on other secondary host plants than crops and maintains the diversity of the species, serving as the source of variability.

In conclusion, the use of genomic approaches has allowed the detection of geographic structure signature in a highly migratory pest species at a national and regional scale. The study has identified two genetic clusters in this aphid pest in England, which could correspond to ecotypes or play different roles as vectors of plant viruses. In addition, there is significant genetic differentiation across the distribution of *R. padi*, being the Southwest population the most differentiated of these. This genetic differentiation cannot be explained just by geographic distance, which suggests that there are other factors that prevent complete panmixia. The dominant form of reproduction in English populations of *R. padi* is mostly sexual, which has an impact on the transmission of BYDV, although population monitoring is still recommended to identify any possible shift towards anholocycly. There is no evidence for long-term demographic changes in the populations of *R. padi*, which is consistent with previous studies of the population dynamics of the species, indicating that environmental change and seasonal selective pressures as insecticide application will have little impact on the genetic diversity of the species. These results have direct implications in control and management of *R. padi*, but further studies are needed to fully understand the diversity and dynamics of the species to improve control programmes and prediction models.

### Data Archiving

MinION (for genome assembly) and Illumina reads (used for population genetics) are available in the European Nucleotide Archive (ENA) database with project numbers PRJEB35176 and PRJEB35188, respectively. The new version of the *R. padi* genome has been uploaded to AphidBase https://bipaa.genouest.org/is/.

## Acknowledgements

This study has been possible thanks to Rothamsted Smart Crop Protection programme and the Rothamsted Insect Survey National Capability grant. The work at Rothamsted forms part of the Smart Crop Protection (SCP) strategic programme (BBS/OS/CP/000001) funded through the Biotechnology and Biological Sciences Research Council’s Industry Strategy Challenge Fund. The Rothamsted Insect Survey, a National Capability, is funded by the Biotechnology and Biological Sciences Research Council under the Core Capability Grant BBS/E/C/000J0200. We would like to thank Peter Thorpe and Jorunn Bos for allowing access to the *Rhopalosiphum padi* genome assembly before publication, providing the trained set of genes and for useful discussions about the genome assembly.

