## supplementary material for "Genetic structure at national and regional scale in a long-distance dispersing pest organism, the bird cherry–oat aphid *Rhopalosiphum padi*"

**Figure S1.** Minion read length frequency histogram.

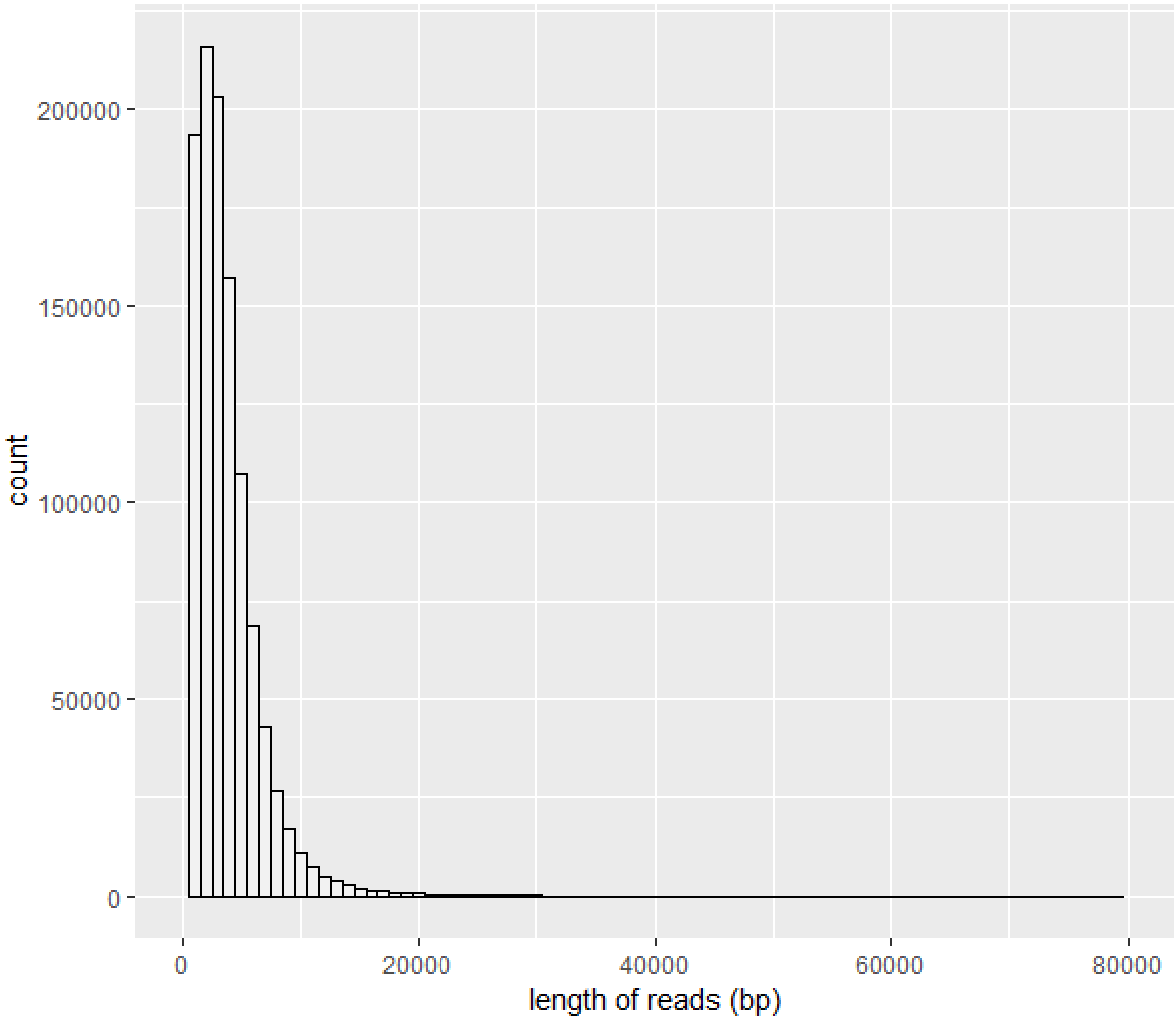

**Figure S2.** BLAST **t**op-hit species distribution.

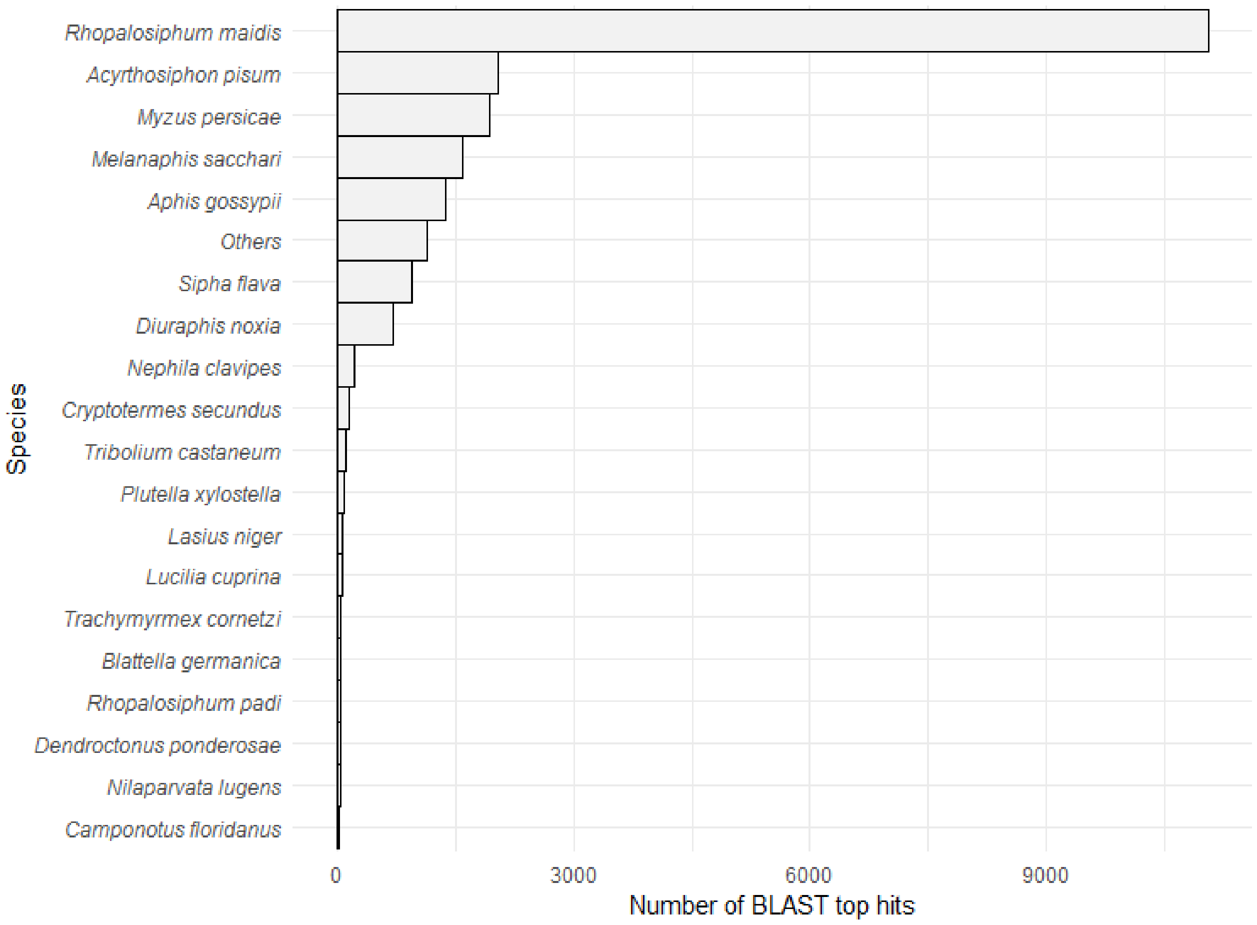

**Figure S3.** Similarity range of the BLAST hits.

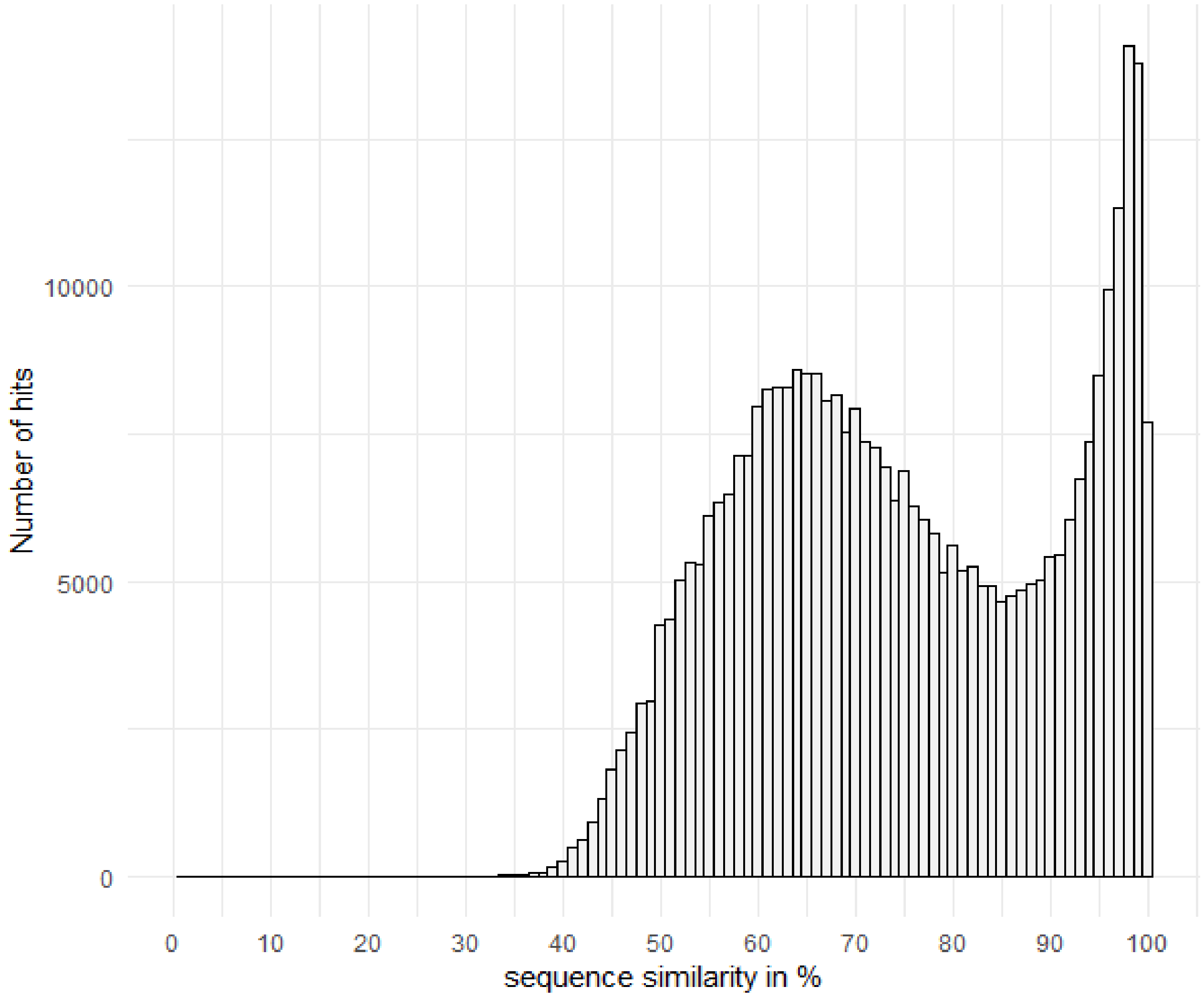

**Figure S4.** Boxplot showing the DNA yield obtained kits from samples collected in different year using different DNA extraction.

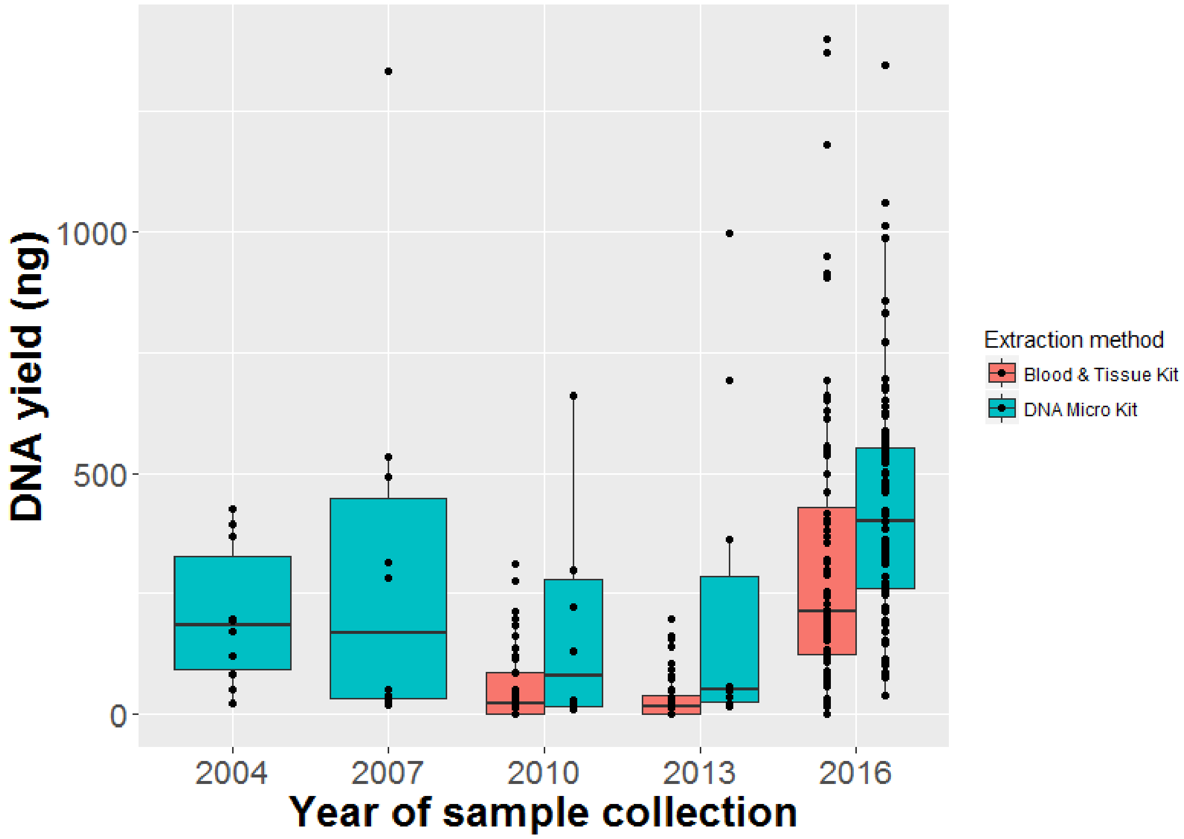

**Figure S5.** Missing data per individual when all samples are analysed together.

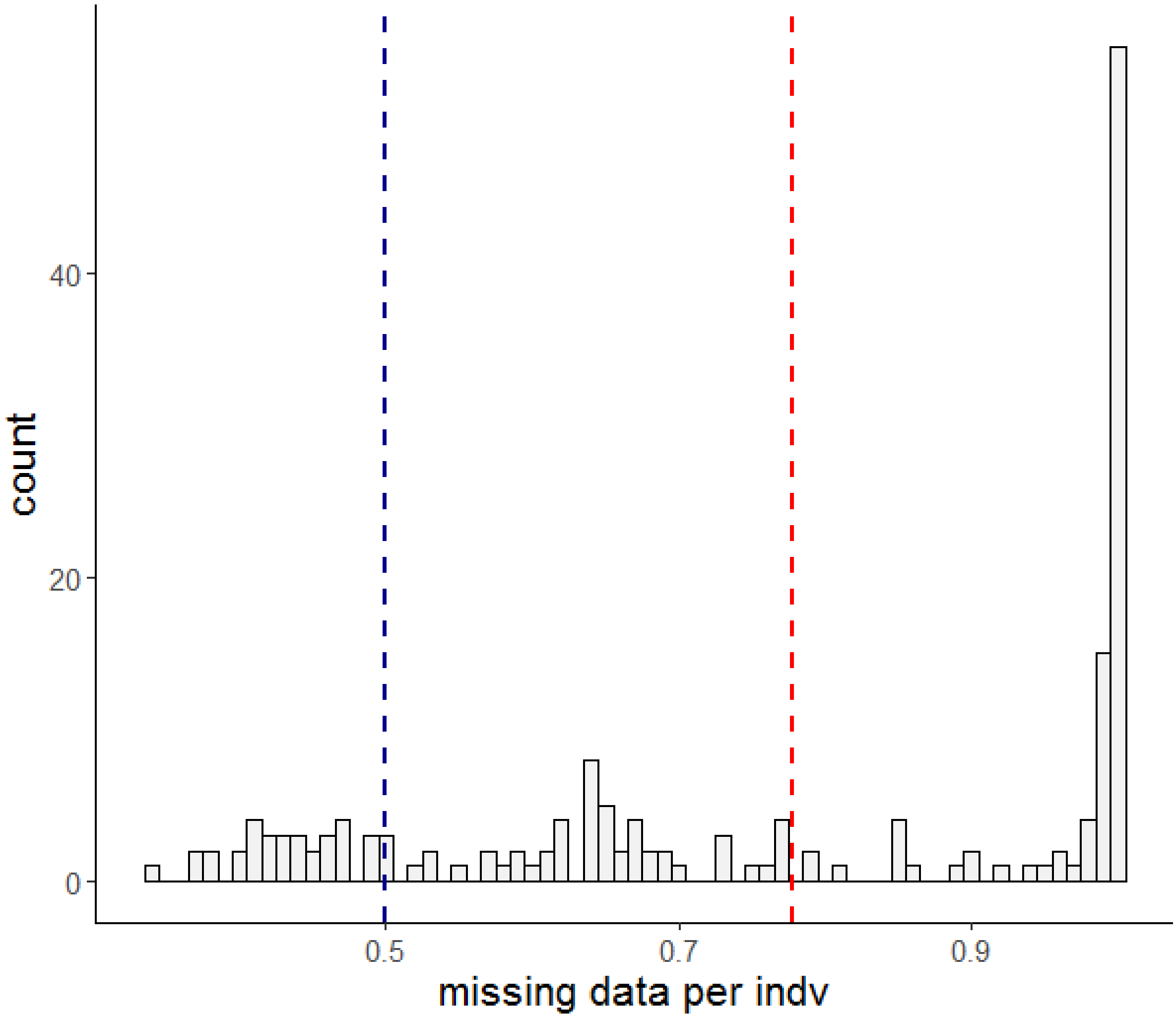

**Figure S6.** Missing data per locus when all samples are analysed together.

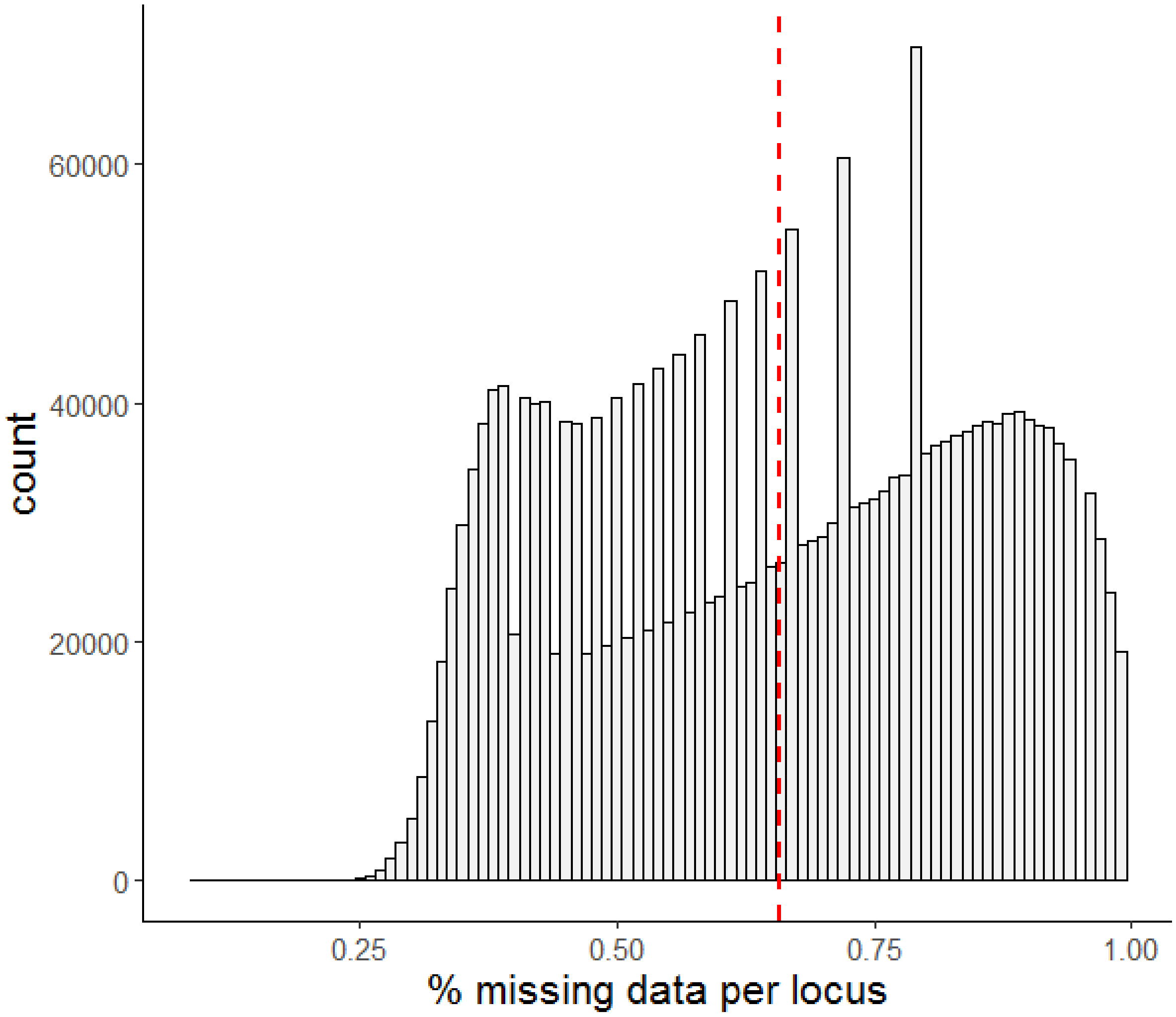

**Figure S7.** PCA analysis of all samples data set (FS7) with no evident clustering according to library or genotyping experiment.

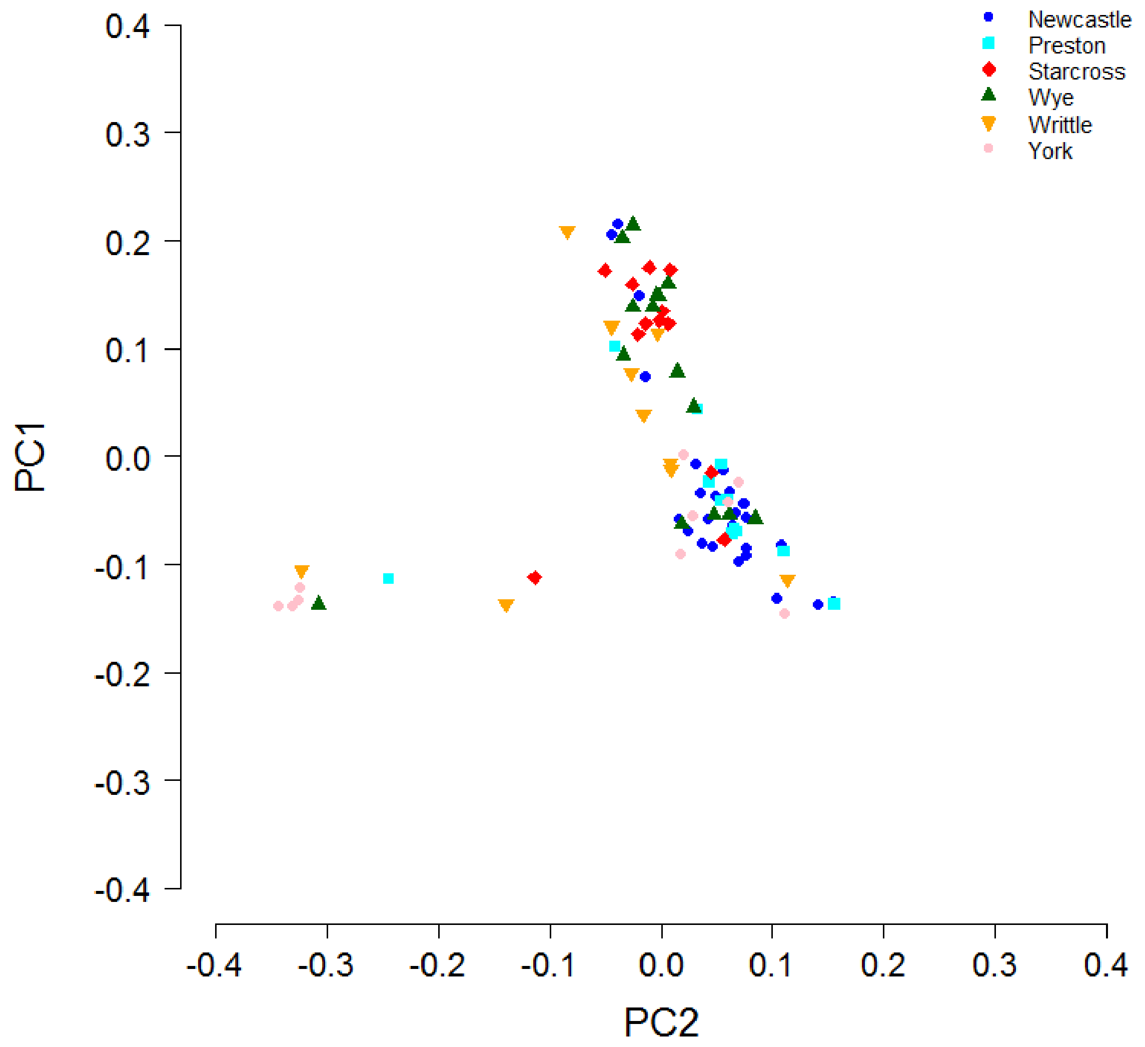

**Figure S8.** Detection of the most likely number of genetic groups (*k*) using Structure and following the statistic Δ(*k*) described in the Evanno *et al*. (2005) method. The likelihood is maximised when *k* = 2.

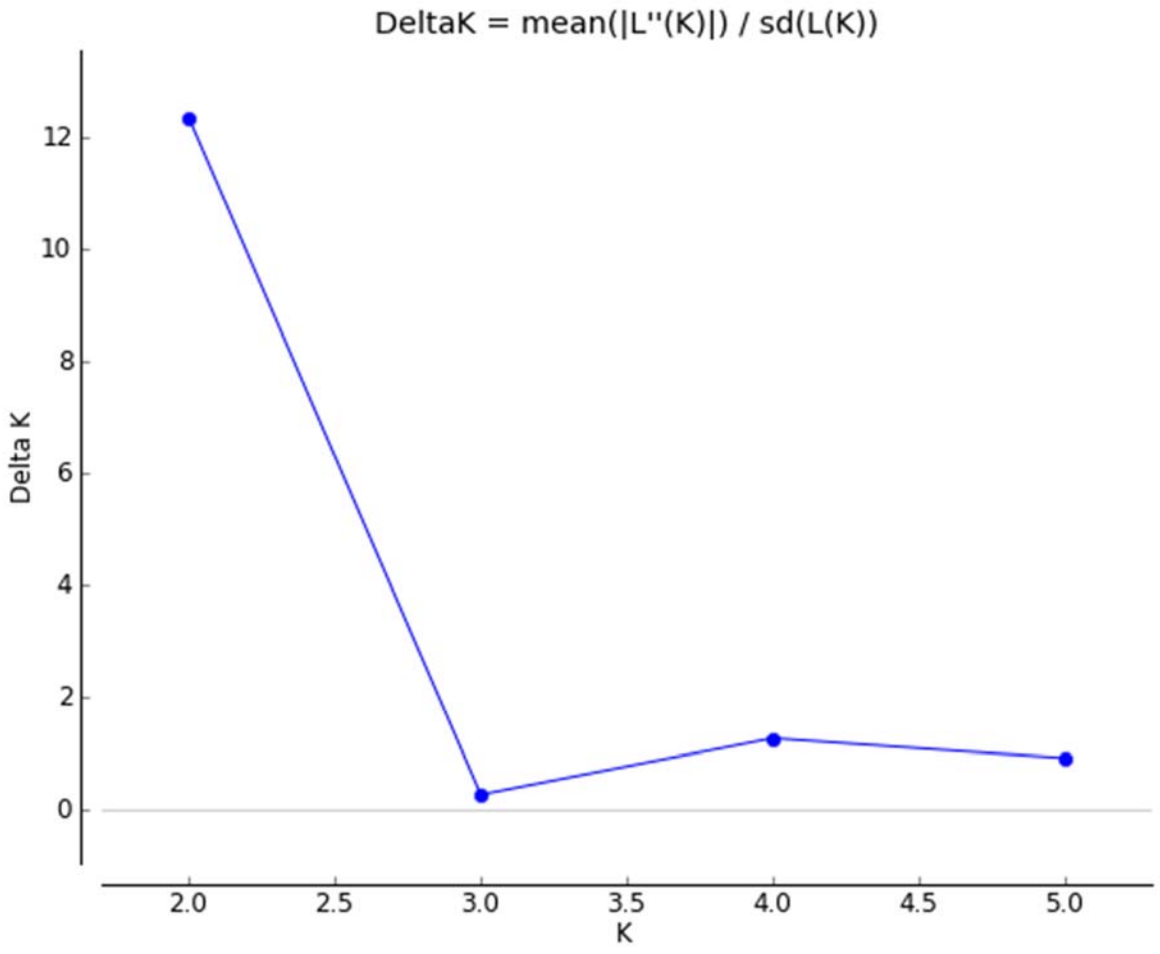

**Figure S9.** Missing data per locus individual after filtering the Newcastle (A and B) and Starcross (C and D) SNP dataset with the FS2. Dashed vertical lines indicate the mean missing data.

**A)**

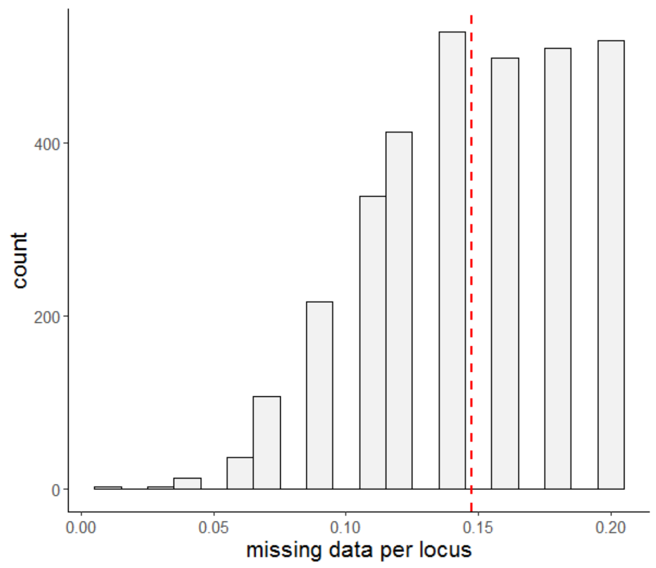


**B)**

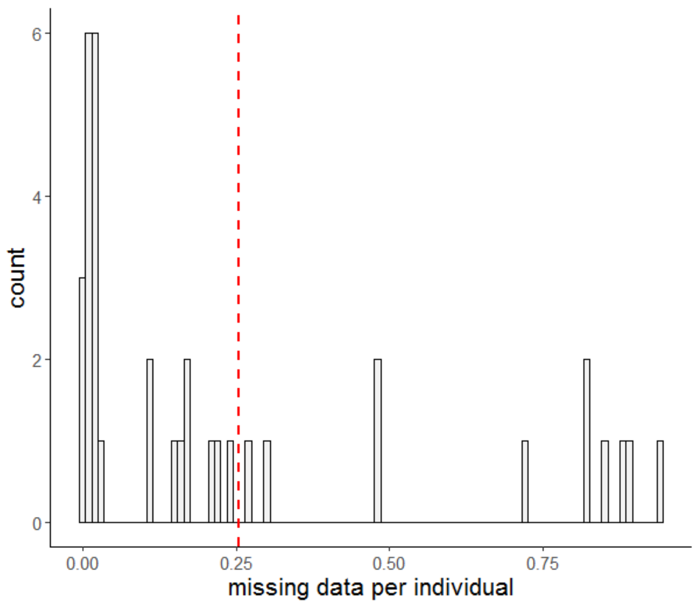


**C)**

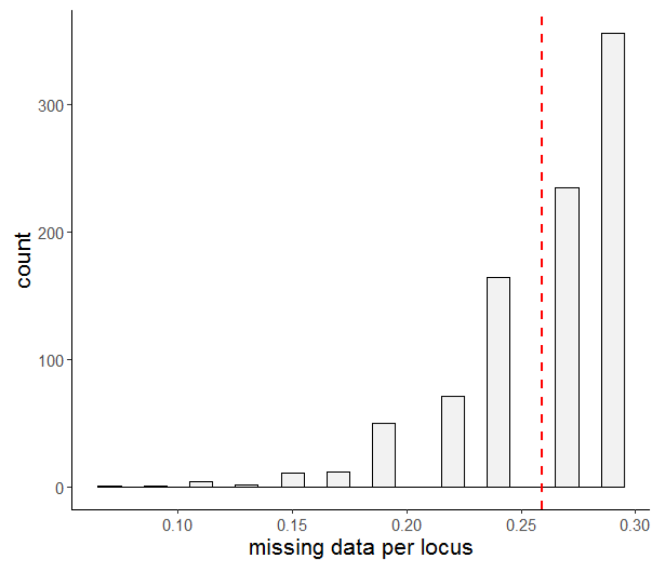


**D)**

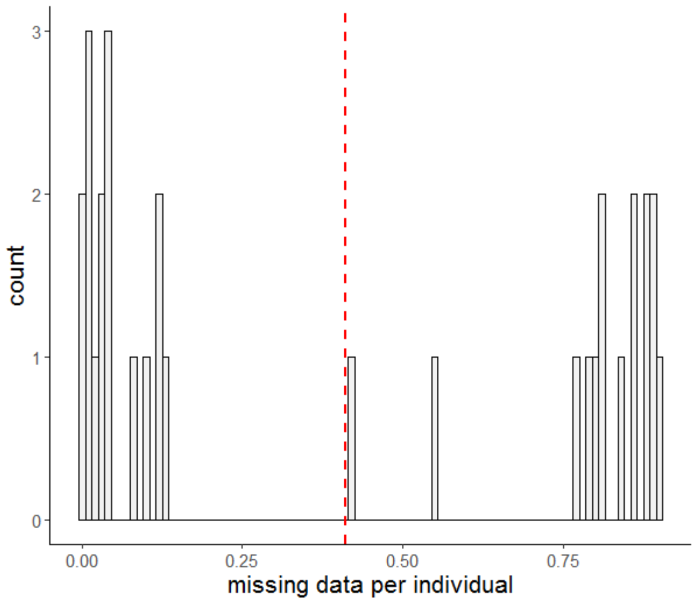

*DNA yield comparison across sample years*

Of the 316 aphids used in the study, 34 samples yielded less than 0.01 ng (Supplementary Table S1). Of these, 19 were samples collected in 2013, 14 in 2010 and 1 aphid individual was from 2016; in addition, all these samples were extracted using Qiagen's Blood & Tissue kit. Aphids from 2016 yielded on average the highest amount of DNA (382.04 ng ± 278.15 standard deviation), followed by samples from 2007 (311.73 ng ± 410.28), 2004 (203.32 ng ± 145.33), 2010 (80.21 ng ± 124.17) and 2013 (70.54 ng ± 168.64). However, the variation in yield from samples within each year is high and they range from 23.82 ng to 426 ng (2004), 18.6 – 1332 ng (2007), 9.09 - 660 ng (2010), 12.1 – 996 ng (2013), and 16.56 – 1400 ng (2016) (Supplementary Table S1). The amount of DNA obtained from aphids from 2010 and 2013 was significantly smaller than that obtained from 2004, 2007 and 2016 samples (Wilcoxon rank sum test $P < 0.01$, after Bonferroni correction), the average DNA obtained from 2016 samples was significantly higher than that from samples collected in other years ($P < 0.005$ after Bonferroni correction) except 2007 (Wilcoxon rank sum test $P = 0.2431$ after Bonferroni correction), and there is no significant difference between the DNA obtained from samples from 2007 and 2004 (Wilcoxon rank sum test $P = 1$ after Bonferroni correction). In addition, the two different DNA extraction kits used in samples from 2010, 2013 and 2016 provided different amounts of DNA on average, with Qiagen's DNA Micro kit resulting in higher amounts (Table 1, Figure S2), and significantly different in the samples from 2010 and 2016 (Wilcoxon rank sum test 2010: $P = 0.024$; 2013: $P = 0.81$; 2016: $P = 7.8 \times 10^{-11}$, with Bonferroni correction).

**Table S1.** Samples used in the present study to extract DNA. Date - when the sample was collected in the suction-trap; Extraction method – the kit used to extract DNA from individual aphids; Conc – concentration as measured using Qubit; Total – the amount of DNA yielded by the specimen.

| Sample | Location | Trap date | Extraction method | Conc (ng/ul) | Total ammount (ng) |
|---|---|---|---|---|---|
| RpH1 | Hereford | 2016 | DNA Micro Kit | 6.47 | 194.1 |
| RpH10 | Hereford | 2016 | DNA Micro Kit | 3.76 | 112.8 |
| RpH11 | Hereford | 2016 | DNA Micro Kit | 13.3 | 399 |
| RpH12 | Hereford | 2016 | DNA Micro Kit | 10.4 | 312 |
| RpH13 | Hereford | 2016 | DNA Micro Kit | 3.8 | 114 |
| RpH14 | Hereford | 2016 | DNA Micro Kit | 22.7 | 681 |
| RpH15 | Hereford | 2016 | DNA Micro Kit | 19 | 570 |
| RpH16 | Hereford | 2016 | DNA Micro Kit | 10.5 | 315 |
| RpH17 | Hereford | 2016 | DNA Micro Kit | 11.7 | 351 |
| RpH18 | Hereford | 2016 | DNA Micro Kit | 44.9 | 1347 |
| RpH19 | Hereford | 2016 | DNA Micro Kit | 19.6 | 588 |
| RpH2 | Hereford | 2016 | DNA Micro Kit | 7.16 | 214.8 |
| RpH20 | Hereford | 2016 | DNA Micro Kit | 20.9 | 627 |
| RpH21 | Hereford | 2016 | DNA Micro Kit | 10.7 | 321 |
| RpH22 | Hereford | 2016 | DNA Micro Kit | 17.5 | 525 |
| RpH23 | Hereford | 2016 | DNA Micro Kit | 27.7 | 831 |
| RpH24 | Hereford | 2016 | DNA Micro Kit | 8.63 | 258.9 |
| RpH25 | Hereford | 2016 | DNA Micro Kit | 5.05 | 151.5 |
| RpH3 | Hereford | 2016 | DNA Micro Kit | 10.7 | 321 |
| RpH4 | Hereford | 2016 | DNA Micro Kit | 23.2 | 696 |
| RpH5 | Hereford | 2016 | DNA Micro Kit | 25.7 | 771 |
| RpH6 | Hereford | 2016 | DNA Micro Kit | 8.44 | 253.2 |
| RpH7 | Hereford | 2016 | DNA Micro Kit | 14.1 | 423 |
| RpH8 | Hereford | 2016 | DNA Micro Kit | 11.6 | 348 |
| RpH9 | Hereford | 2016 | DNA Micro Kit | 9.16 | 274.8 |
| RpN36 | Newcastle | 2004 | DNA Micro Kit | 12.3 | 369 |
| RpN37 | Newcastle | 2004 | DNA Micro Kit | 1.75 | 52.5 |
| RpN38 | Newcastle | 2004 | DNA Micro Kit | 14.2 | 426 |
| RpN39 | Newcastle | 2004 | DNA Micro Kit | 13.1 | 393 |
| RpN40 | Newcastle | 2004 | DNA Micro Kit | 6.47 | 194.1 |
| RpN61 | Newcastle | 2004 | DNA Micro Kit | 13.6 | 408 |
| RpN62 | Newcastle | 2004 | DNA Micro Kit | 5.55 | 166.5 |
| RpN63 | Newcastle | 2004 | DNA Micro Kit | 4.6 | 138 |
| RpN64 | Newcastle | 2004 | DNA Micro Kit | 6.66 | 199.8 |
| RpN65 | Newcastle | 2004 | DNA Micro Kit | 3.64 | 109.2 |
| RpN51 | Newcastle | 2007 | DNA Micro Kit | 17.8 | 534 |
| RpN52 | Newcastle | 2007 | DNA Micro Kit | 10.5 | 315 |
| RpN53 | Newcastle | 2007 | DNA Micro Kit | 16.4 | 492 |
| RpN54 | Newcastle | 2007 | DNA Micro Kit | 44.4 | 1332 |
| RpN55 | Newcastle | 2007 | DNA Micro Kit | 9.43 | 282.9 |
| RpN66 | Newcastle | 2007 | DNA Micro Kit | 7.09 | 212.7 |
| RpN67 | Newcastle | 2007 | DNA Micro Kit | 9.65 | 289.5 |

| | | | | | |
|---|---|---|---|---|---|
| RpN68 | Newcastle | 2007 | DNA Micro Kit | 23.8 | 714 |
| RpN69 | Newcastle | 2007 | DNA Micro Kit | 19.2 | 576 |
| RpN70 | Newcastle | 2007 | DNA Micro Kit | 17.6 | 528 |
| RpN26 | Newcastle | 2010 | Blood & Tissue Kit | 1.9 | 47.5 |
| RpN27 | Newcastle | 2010 | Blood & Tissue Kit | 7.36 | 184 |
| RpN28 | Newcastle | 2010 | Blood & Tissue Kit | 5.48 | 137 |
| RpN29 | Newcastle | 2010 | Blood & Tissue Kit | 8.56 | 214 |
| RpN30 | Newcastle | 2010 | Blood & Tissue Kit | 4.64 | 116 |
| RpN31 | Newcastle | 2010 | Blood & Tissue Kit | 6.48 | 162 |
| RpN32 | Newcastle | 2010 | Blood & Tissue Kit | 1.3 | 32.5 |
| RpN33 | Newcastle | 2010 | Blood & Tissue Kit | 11.1 | 277.5 |
| RpN34 | Newcastle | 2010 | Blood & Tissue Kit | 12.5 | 312.5 |
| RpN35 | Newcastle | 2010 | Blood & Tissue Kit | 7.88 | 197 |
| RpN46 | Newcastle | 2010 | DNA Micro Kit | 4.36 | 130.8 |
| RpN47 | Newcastle | 2010 | DNA Micro Kit | 9.97 | 299.1 |
| RpN48 | Newcastle | 2010 | DNA Micro Kit | 7.4 | 222 |
| RpN49 | Newcastle | 2010 | DNA Micro Kit | 9.93 | 297.9 |
| RpN50 | Newcastle | 2010 | DNA Micro Kit | 22 | 660 |
| RpN71 | Newcastle | 2010 | DNA Micro Kit | 2.89 | 86.7 |
| RpN72 | Newcastle | 2010 | DNA Micro Kit | 9.54 | 286.2 |
| RpN73 | Newcastle | 2010 | DNA Micro Kit | 7.55 | 226.5 |
| RpN74 | Newcastle | 2010 | DNA Micro Kit | 3.74 | 112.2 |
| RpN75 | Newcastle | 2010 | DNA Micro Kit | 16.2 | 486 |
| RpN16 | Newcastle | 2013 | Blood & Tissue Kit | 0.492 | 12.3 |
| RpN17 | Newcastle | 2013 | Blood & Tissue Kit | 7.92 | 198 |
| RpN18 | Newcastle | 2013 | Blood & Tissue Kit | 0.788 | 19.7 |
| RpN19 | Newcastle | 2013 | Blood & Tissue Kit | 6.28 | 157 |
| RpN20 | Newcastle | 2013 | Blood & Tissue Kit | 3.69 | 92.25 |
| RpN21 | Newcastle | 2013 | Blood & Tissue Kit | 0.732 | 18.3 |
| RpN22 | Newcastle | 2013 | Blood & Tissue Kit | 2 | 50 |
| RpN23 | Newcastle | 2013 | Blood & Tissue Kit | 1.98 | 49.5 |
| RpN24 | Newcastle | 2013 | Blood & Tissue Kit | 6.56 | 164 |
| RpN25 | Newcastle | 2013 | Blood & Tissue Kit | 0.836 | 20.9 |
| RpN56 | Newcastle | 2013 | DNA Micro Kit | 0.696 | 20.88 |
| RpN57 | Newcastle | 2013 | DNA Micro Kit | 1.21 | 36.3 |
| RpN58 | Newcastle | 2013 | DNA Micro Kit | 33.2 | 996 |
| RpN59 | Newcastle | 2013 | DNA Micro Kit | 23.1 | 693 |
| RpN60 | Newcastle | 2013 | DNA Micro Kit | 12.1 | 363 |
| RpN76 | Newcastle | 2013 | DNA Micro Kit | 6.6 | 198 |
| RpN77 | Newcastle | 2013 | DNA Micro Kit | 1.81 | 54.3 |
| RpN78 | Newcastle | 2013 | DNA Micro Kit | 12.2 | 366 |
| RpN79 | Newcastle | 2013 | DNA Micro Kit | 4.61 | 138.3 |
| RpN80 | Newcastle | 2013 | DNA Micro Kit | 2.62 | 78.6 |
| RpN1 | Newcastle | 2016 | Blood & Tissue Kit | 38 | 950 |
| RpN10 | Newcastle | 2016 | Blood & Tissue Kit | 20 | 500 |
| RpN11 | Newcastle | 2016 | Blood & Tissue Kit | 10.2 | 255 |

| | | | | | |
|---|---|---|---|---|---|
| RpN12 | Newcastle | 2016 | Blood & Tissue Kit | 9.8 | 245 |
| RpN13 | Newcastle | 2016 | Blood & Tissue Kit | 27.7 | 692.5 |
| RpN14 | Newcastle | 2016 | Blood & Tissue Kit | 19.9 | 497.5 |
| RpN15 | Newcastle | 2016 | Blood & Tissue Kit | 7.44 | 186 |
| RpN2 | Newcastle | 2016 | Blood & Tissue Kit | 47.2 | 1180 |
| RpN3 | Newcastle | 2016 | Blood & Tissue Kit | 36.2 | 905 |
| RpN4 | Newcastle | 2016 | Blood & Tissue Kit | 36.6 | 915 |
| RpN5 | Newcastle | 2016 | Blood & Tissue Kit | 54.8 | 1370 |
| RpN6 | Newcastle | 2016 | Blood & Tissue Kit | 26 | 650 |
| RpN7 | Newcastle | 2016 | Blood & Tissue Kit | 22.2 | 555 |
| RpN8 | Newcastle | 2016 | Blood & Tissue Kit | 26.4 | 660 |
| RpN9 | Newcastle | 2016 | Blood & Tissue Kit | 56 | 1400 |
| RpN41 | Newcastle | 2016 | DNA Micro Kit | 15.8 | 474 |
| RpN42 | Newcastle | 2016 | DNA Micro Kit | 9.16 | 274.8 |
| RpN43 | Newcastle | 2016 | DNA Micro Kit | 32.9 | 987 |
| RpN44 | Newcastle | 2016 | DNA Micro Kit | 8.46 | 253.8 |
| RpN45 | Newcastle | 2016 | DNA Micro Kit | 22.4 | 672 |
| RpP27 | Preston | 2010 | Blood & Tissue Kit | 3.52 | 88 |
| RpP28 | Preston | 2010 | Blood & Tissue Kit | 0.58 | 14.5 |
| RpP29 | Preston | 2010 | Blood & Tissue Kit | 4.88 | 122 |
| RpP30 | Preston | 2010 | Blood & Tissue Kit | <0.01 | 0 |
| RpP31 | Preston | 2010 | Blood & Tissue Kit | <0.01 | 0 |
| RpP32 | Preston | 2010 | Blood & Tissue Kit | 3.41 | 85.25 |
| RpP33 | Preston | 2010 | Blood & Tissue Kit | <0.01 | 0 |
| RpP34 | Preston | 2010 | Blood & Tissue Kit | 1.09 | 27.25 |
| RpP35 | Preston | 2010 | Blood & Tissue Kit | 0.952 | 23.8 |
| RpP36 | Preston | 2010 | Blood & Tissue Kit | 1.45 | 36.25 |
| RpP17 | Preston | 2013 | Blood & Tissue Kit | 0.992 | 24.8 |
| RpP18 | Preston | 2013 | Blood & Tissue Kit | <0.01 | 0 |
| RpP19 | Preston | 2013 | Blood & Tissue Kit | <0.01 | 0 |
| RpP20 | Preston | 2013 | Blood & Tissue Kit | 0.828 | 20.7 |
| RpP21 | Preston | 2013 | Blood & Tissue Kit | 1.49 | 37.25 |
| RpP22 | Preston | 2013 | Blood & Tissue Kit | 3.1 | 77.5 |
| RpP23 | Preston | 2013 | Blood & Tissue Kit | <0.01 | 0 |
| RpP24 | Preston | 2013 | Blood & Tissue Kit | 0.892 | 22.3 |
| RpP25 | Preston | 2013 | Blood & Tissue Kit | 0.512 | 12.8 |
| RpP26 | Preston | 2013 | Blood & Tissue Kit | <0.01 | 0 |
| RpP37 | Preston | 2013 | Blood & Tissue Kit | 1.04 | 26 |
| RpP38 | Preston | 2013 | Blood & Tissue Kit | <0.01 | 0 |
| RpP39 | Preston | 2013 | Blood & Tissue Kit | 4.24 | 106 |
| RpP40 | Preston | 2013 | Blood & Tissue Kit | 3.22 | 80.5 |
| RpP1 | Preston | 2016 | Blood & Tissue Kit | 7.08 | 177 |
| RpP10 | Preston | 2016 | Blood & Tissue Kit | 7.36 | 184 |
| RpP11 | Preston | 2016 | Blood & Tissue Kit | 11.9 | 297.5 |
| RpP12 | Preston | 2016 | Blood & Tissue Kit | 21.5 | 537.5 |
| RpP13 | Preston | 2016 | Blood & Tissue Kit | 8.16 | 204 |

| | | | | | |
|---|---|---|---|---|---|
| RpP14 | Preston | 2016 | Blood & Tissue Kit | 11.6 | 290 |
| RpP15 | Preston | 2016 | Blood & Tissue Kit | 12.9 | 322.5 |
| RpP16 | Preston | 2016 | Blood & Tissue Kit | 8.64 | 216 |
| RpP2 | Preston | 2016 | Blood & Tissue Kit | 6.96 | 174 |
| RpP3 | Preston | 2016 | Blood & Tissue Kit | 25.1 | 627.5 |
| RpP4 | Preston | 2016 | Blood & Tissue Kit | 3.43 | 85.75 |
| RpP5 | Preston | 2016 | Blood & Tissue Kit | 24.5 | 612.5 |
| RpP6 | Preston | 2016 | Blood & Tissue Kit | 3.19 | 79.75 |
| RpP7 | Preston | 2016 | Blood & Tissue Kit | 7.04 | 176 |
| RpP8 | Preston | 2016 | Blood & Tissue Kit | 14.8 | 370 |
| RpP9 | Preston | 2016 | Blood & Tissue Kit | 4.76 | 119 |
| RpSX67 | Starcross | 2004 | DNA Micro Kit | 4.09 | 122.7 |
| RpSX68 | Starcross | 2004 | DNA Micro Kit | 0.794 | 23.82 |
| RpSX69 | Starcross | 2004 | DNA Micro Kit | 5.75 | 172.5 |
| RpSX70 | Starcross | 2004 | DNA Micro Kit | 6.56 | 196.8 |
| RpSX71 | Starcross | 2004 | DNA Micro Kit | 2.76 | 82.8 |
| RpSX87 | Starcross | 2004 | DNA Micro Kit | 6.01 | 180.3 |
| RpSX88 | Starcross | 2004 | DNA Micro Kit | 0.58 | 17.4 |
| RpSX89 | Starcross | 2004 | DNA Micro Kit | 1.83 | 54.9 |
| RpSX90 | Starcross | 2004 | DNA Micro Kit | 6.09 | 182.7 |
| RpSX91 | Starcross | 2004 | DNA Micro Kit | 5.33 | 159.9 |
| RpSX77 | Starcross | 2007 | DNA Micro Kit | 1.76 | 52.8 |
| RpSX78 | Starcross | 2007 | DNA Micro Kit | 0.62 | 18.6 |
| RpSX79 | Starcross | 2007 | DNA Micro Kit | 0.973 | 29.19 |
| RpSX80 | Starcross | 2007 | DNA Micro Kit | 0.697 | 20.91 |
| RpSX81 | Starcross | 2007 | DNA Micro Kit | 1.33 | 39.9 |
| RpSX92 | Starcross | 2007 | DNA Micro Kit | 1.27 | 38.1 |
| RpSX93 | Starcross | 2007 | DNA Micro Kit | 1.12 | 33.6 |
| RpSX94 | Starcross | 2007 | DNA Micro Kit | 1.33 | 39.9 |
| RpSX95 | Starcross | 2007 | DNA Micro Kit | 1.25 | 37.5 |
| RpSX96 | Starcross | 2007 | DNA Micro Kit | 1.03 | 30.9 |
| RpSX26 | Starcross | 2010 | Blood & Tissue Kit | 0.46 | 23 |
| RpSX27 | Starcross | 2010 | Blood & Tissue Kit | 0.548 | 27.4 |
| RpSX28 | Starcross | 2010 | Blood & Tissue Kit | <0.01 | 0 |
| RpSX29 | Starcross | 2010 | Blood & Tissue Kit | <0.01 | 0 |
| RpSX30 | Starcross | 2010 | Blood & Tissue Kit | <0.01 | 0 |
| RpSX31 | Starcross | 2010 | Blood & Tissue Kit | 0.852 | 21.3 |
| RpSX32 | Starcross | 2010 | Blood & Tissue Kit | 0.684 | 17.1 |
| RpSX33 | Starcross | 2010 | Blood & Tissue Kit | <0.01 | 0 |
| RpSX34 | Starcross | 2010 | Blood & Tissue Kit | 2.02 | 50.5 |
| RpSX35 | Starcross | 2010 | Blood & Tissue Kit | 0.788 | 19.7 |
| RpSX57 | Starcross | 2010 | Blood & Tissue Kit | <0.01 | 0 |
| RpSX58 | Starcross | 2010 | Blood & Tissue Kit | <0.01 | 0 |
| RpSX59 | Starcross | 2010 | Blood & Tissue Kit | 0.556 | 13.9 |
| RpSX60 | Starcross | 2010 | Blood & Tissue Kit | <0.01 | 0 |
| RpSX61 | Starcross | 2010 | Blood & Tissue Kit | <0.01 | 0 |

| | | | | | |
|---|---|---|---|---|---|
| RpSX62 | Starcross | 2010 | Blood & Tissue Kit | 0.716 | 17.9 |
| RpSX63 | Starcross | 2010 | Blood & Tissue Kit | <0.01 | 0 |
| RpSX64 | Starcross | 2010 | Blood & Tissue Kit | 1.86 | 46.5 |
| RpSX65 | Starcross | 2010 | Blood & Tissue Kit | <0.01 | 0 |
| RpSX66 | Starcross | 2010 | Blood & Tissue Kit | <0.01 | 0 |
| RpSX100 | Starcross | 2010 | DNA Micro Kit | 0.516 | 15.48 |
| RpSX101 | Starcross | 2010 | DNA Micro Kit | 0.695 | 20.85 |
| RpSX82 | Starcross | 2010 | DNA Micro Kit | 0.303 | 9.09 |
| RpSX83 | Starcross | 2010 | DNA Micro Kit | 0.446 | 13.38 |
| RpSX84 | Starcross | 2010 | DNA Micro Kit | 0.455 | 13.65 |
| RpSX85 | Starcross | 2010 | DNA Micro Kit | 0.958 | 28.74 |
| RpSX86 | Starcross | 2010 | DNA Micro Kit | 0.721 | 21.63 |
| RpSX97 | Starcross | 2010 | DNA Micro Kit | 0.692 | 20.76 |
| RpSX98 | Starcross | 2010 | DNA Micro Kit | 0.817 | 24.51 |
| RpSX99 | Starcross | 2010 | DNA Micro Kit | 0.674 | 20.22 |
| RpSX16 | Starcross | 2013 | Blood & Tissue Kit | <0.01 | 0 |
| RpSX17 | Starcross | 2013 | Blood & Tissue Kit | 0.984 | 19.68 |
| RpSX18 | Starcross | 2013 | Blood & Tissue Kit | <0.01 | 0 |
| RpSX19 | Starcross | 2013 | Blood & Tissue Kit | <0.01 | 0 |
| RpSX20 | Starcross | 2013 | Blood & Tissue Kit | <0.01 | 0 |
| RpSX21 | Starcross | 2013 | Blood & Tissue Kit | <0.01 | 0 |
| RpSX22 | Starcross | 2013 | Blood & Tissue Kit | <0.01 | 0 |
| RpSX23 | Starcross | 2013 | Blood & Tissue Kit | <0.01 | 0 |
| RpSX24 | Starcross | 2013 | Blood & Tissue Kit | 0.608 | 30.4 |
| RpSX25 | Starcross | 2013 | Blood & Tissue Kit | <0.01 | 0 |
| RpSX47 | Starcross | 2013 | Blood & Tissue Kit | <0.01 | 0 |
| RpSX48 | Starcross | 2013 | Blood & Tissue Kit | <0.01 | 0 |
| RpSX49 | Starcross | 2013 | Blood & Tissue Kit | <0.01 | 0 |
| RpSX50 | Starcross | 2013 | Blood & Tissue Kit | <0.01 | 0 |
| RpSX51 | Starcross | 2013 | Blood & Tissue Kit | <0.01 | 0 |
| RpSX52 | Starcross | 2013 | Blood & Tissue Kit | 5.64 | 141 |
| RpSX53 | Starcross | 2013 | Blood & Tissue Kit | <0.01 | 0 |
| RpSX54 | Starcross | 2013 | Blood & Tissue Kit | 1.45 | 36.25 |
| RpSX55 | Starcross | 2013 | Blood & Tissue Kit | 3 | 75 |
| RpSX56 | Starcross | 2013 | Blood & Tissue Kit | 0.484 | 12.1 |
| RpSX102 | Starcross | 2013 | DNA Micro Kit | 0.535 | 16.05 |
| RpSX103 | Starcross | 2013 | DNA Micro Kit | 0.37 | 11.1 |
| RpSX104 | Starcross | 2013 | DNA Micro Kit | 0.419 | 12.57 |
| RpSX105 | Starcross | 2013 | DNA Micro Kit | 0.624 | 18.72 |
| RpSX106 | Starcross | 2013 | DNA Micro Kit | 0.419 | 12.57 |
| RpSX72 | Starcross | 2013 | DNA Micro Kit | 1.88 | 56.4 |
| RpSX73 | Starcross | 2013 | DNA Micro Kit | 1.66 | 49.8 |
| RpSX74 | Starcross | 2013 | DNA Micro Kit | 0.722 | 21.66 |
| RpSX75 | Starcross | 2013 | DNA Micro Kit | 1.67 | 50.1 |
| RpSX76 | Starcross | 2013 | DNA Micro Kit | 0.595 | 17.85 |
| RpSX1 | Starcross | 2016 | Blood & Tissue Kit | 4.54 | 90.8 |

| | | | | | |
|---|---|---|---|---|---|
| RpSX10 | Starcross | 2016 | Blood & Tissue Kit | 6.36 | 127.2 |
| RpSX11 | Starcross | 2016 | Blood & Tissue Kit | 2.95 | 59 |
| RpSX12 | Starcross | 2016 | Blood & Tissue Kit | 5.68 | 113.6 |
| RpSX13 | Starcross | 2016 | Blood & Tissue Kit | 4.03 | 80.6 |
| RpSX14 | Starcross | 2016 | Blood & Tissue Kit | 6.72 | 134.4 |
| RpSX15 | Starcross | 2016 | Blood & Tissue Kit | 5.88 | 117.6 |
| RpSX2 | Starcross | 2016 | Blood & Tissue Kit | 0.828 | 16.56 |
| RpSX3 | Starcross | 2016 | Blood & Tissue Kit | 7.92 | 158.4 |
| RpSX36 | Starcross | 2016 | Blood & Tissue Kit | <0.01 | 0 |
| RpSX37 | Starcross | 2016 | Blood & Tissue Kit | 6.72 | 168 |
| RpSX38 | Starcross | 2016 | Blood & Tissue Kit | 15.2 | 380 |
| RpSX39 | Starcross | 2016 | Blood & Tissue Kit | 6.32 | 158 |
| RpSX4 | Starcross | 2016 | Blood & Tissue Kit | 17.8 | 356 |
| RpSX40 | Starcross | 2016 | Blood & Tissue Kit | 1.08 | 27 |
| RpSX41 | Starcross | 2016 | Blood & Tissue Kit | 5.2 | 130 |
| RpSX42 | Starcross | 2016 | Blood & Tissue Kit | 2.93 | 73.25 |
| RpSX43 | Starcross | 2016 | Blood & Tissue Kit | 1.3 | 32.5 |
| RpSX44 | Starcross | 2016 | Blood & Tissue Kit | 3.22 | 80.5 |
| RpSX45 | Starcross | 2016 | Blood & Tissue Kit | 6.96 | 174 |
| RpSX46 | Starcross | 2016 | Blood & Tissue Kit | 8.08 | 202 |
| RpSX5 | Starcross | 2016 | Blood & Tissue Kit | 11.4 | 228 |
| RpSX6 | Starcross | 2016 | Blood & Tissue Kit | 3.44 | 68.8 |
| RpSX7 | Starcross | 2016 | Blood & Tissue Kit | 5.44 | 108.8 |
| RpSX8 | Starcross | 2016 | Blood & Tissue Kit | 4.4 | 88 |
| RpSX9 | Starcross | 2016 | Blood & Tissue Kit | 1.04 | 20.8 |
| RpWr1 | Writtle | 2016 | DNA Micro Kit | 7.44 | 223.2 |
| RpWr10 | Writtle | 2016 | DNA Micro Kit | 8.72 | 261.6 |
| RpWr11 | Writtle | 2016 | DNA Micro Kit | 12.1 | 363 |
| RpWr12 | Writtle | 2016 | DNA Micro Kit | 25.7 | 771 |
| RpWr13 | Writtle | 2016 | DNA Micro Kit | 18.4 | 552 |
| RpWr14 | Writtle | 2016 | DNA Micro Kit | 20.6 | 618 |
| RpWr15 | Writtle | 2016 | DNA Micro Kit | 28.6 | 858 |
| RpWr16 | Writtle | 2016 | DNA Micro Kit | 17.4 | 522 |
| RpWr17 | Writtle | 2016 | DNA Micro Kit | 11.3 | 339 |
| RpWr18 | Writtle | 2016 | DNA Micro Kit | 4.86 | 145.8 |
| RpWr19 | Writtle | 2016 | DNA Micro Kit | 18.7 | 561 |
| RpWr2 | Writtle | 2016 | DNA Micro Kit | 9.51 | 285.3 |
| RpWr20 | Writtle | 2016 | DNA Micro Kit | 14 | 420 |
| RpWr21 | Writtle | 2016 | DNA Micro Kit | 12.8 | 384 |
| RpWr22 | Writtle | 2016 | DNA Micro Kit | 16.7 | 501 |
| RpWr23 | Writtle | 2016 | DNA Micro Kit | 21.3 | 639 |
| RpWr3 | Writtle | 2016 | DNA Micro Kit | 17.9 | 537 |
| RpWr4 | Writtle | 2016 | DNA Micro Kit | 8.74 | 262.2 |
| RpWr5 | Writtle | 2016 | DNA Micro Kit | 3.43 | 102.9 |
| RpWr6 | Writtle | 2016 | DNA Micro Kit | 5.7 | 171 |
| RpWr7 | Writtle | 2016 | DNA Micro Kit | 8.97 | 269.1 |

| | | | | | |
|---|---|---|---|---|---|
| RpWr8 | Writtle | 2016 | DNA Micro Kit | 3.42 | 102.6 |
| RpWr9 | Writtle | 2016 | DNA Micro Kit | 14.1 | 423 |
| RpW1 | Wye | 2016 | Blood & Tissue Kit | 16.1 | 402.5 |
| RpW10 | Wye | 2016 | Blood & Tissue Kit | 18.4 | 460 |
| RpW11 | Wye | 2016 | Blood & Tissue Kit | 18.4 | 460 |
| RpW12 | Wye | 2016 | Blood & Tissue Kit | 16.7 | 417.5 |
| RpW13 | Wye | 2016 | Blood & Tissue Kit | 15.9 | 397.5 |
| RpW14 | Wye | 2016 | Blood & Tissue Kit | 12.6 | 315 |
| RpW15 | Wye | 2016 | Blood & Tissue Kit | 8.68 | 217 |
| RpW2 | Wye | 2016 | Blood & Tissue Kit | 10.2 | 255 |
| RpW3 | Wye | 2016 | Blood & Tissue Kit | 7.68 | 192 |
| RpW4 | Wye | 2016 | Blood & Tissue Kit | 9.92 | 248 |
| RpW5 | Wye | 2016 | Blood & Tissue Kit | 21.8 | 545 |
| RpW6 | Wye | 2016 | Blood & Tissue Kit | 5 | 125 |
| RpW7 | Wye | 2016 | Blood & Tissue Kit | 8.4 | 210 |
| RpW8 | Wye | 2016 | Blood & Tissue Kit | 6.1 | 152.5 |
| RpW9 | Wye | 2016 | Blood & Tissue Kit | 16.1 | 402.5 |
| RpY1 | York | 2016 | DNA Micro Kit | 13.4 | 402 |
| RpY10 | York | 2016 | DNA Micro Kit | 14 | 420 |
| RpY11 | York | 2016 | DNA Micro Kit | 18.2 | 546 |
| RpY12 | York | 2016 | DNA Micro Kit | 15.7 | 471 |
| RpY13 | York | 2016 | DNA Micro Kit | 11.9 | 357 |
| RpY14 | York | 2016 | DNA Micro Kit | 19.3 | 579 |
| RpY15 | York | 2016 | DNA Micro Kit | 11 | 330 |
| RpY16 | York | 2016 | DNA Micro Kit | 19.6 | 588 |
| RpY17 | York | 2016 | DNA Micro Kit | 13.8 | 414 |
| RpY18 | York | 2016 | DNA Micro Kit | 10.6 | 318 |
| RpY19 | York | 2016 | DNA Micro Kit | 11.8 | 354 |
| RpY2 | York | 2016 | DNA Micro Kit | 6.27 | 188.1 |
| RpY20 | York | 2016 | DNA Micro Kit | 8.3 | 249 |
| RpY21 | York | 2016 | DNA Micro Kit | 35.3 | 1059 |
| RpY22 | York | 2016 | DNA Micro Kit | 17.6 | 528 |
| RpY23 | York | 2016 | DNA Micro Kit | 2.53 | 75.9 |
| RpY24 | York | 2016 | DNA Micro Kit | 2.83 | 84.9 |
| RpY25 | York | 2016 | DNA Micro Kit | 12.8 | 384 |
| RpY26 | York | 2016 | DNA Micro Kit | 16.6 | 498 |
| RpY27 | York | 2016 | DNA Micro Kit | 33.8 | 1014 |
| RpY3 | York | 2016 | DNA Micro Kit | 1.27 | 38.1 |
| RpY4 | York | 2016 | DNA Micro Kit | 15.4 | 462 |
| RpY5 | York | 2016 | DNA Micro Kit | 14.1 | 423 |
| RpY6 | York | 2016 | DNA Micro Kit | 21.7 | 651 |
| RpY7 | York | 2016 | DNA Micro Kit | 8.5 | 255 |
| RpY8 | York | 2016 | DNA Micro Kit | 16.6 | 498 |
| RpY9 | York | 2016 | DNA Micro Kit | 16.1 | 483 |

**Table S2.** Details of the samples used for genotype for sequencing (GBS), including the DNA concentration ([DNA]) and amount before and after whole genome amplification (WGA). 2nd WGA refers to the reactions that used the first WGA product as template.

| Sample | Trap date | [DNA] (ng/µl) | Ammount | WGA (ng/µl) | Ammount (ng) | 2nd WGA (ng/µl) | Ammount (ng) | DNA sequenced (ng) |
|---|---|---|---|---|---|---|---|---|
| RpN1 | May 2016 | 38 | 950 | 173 | 3,460 | | | 800 |
| RpN2 | May 2016 | 47.2 | 1180 | 171 | 3,420 | | | 800 |
| RpN3 | May 2016 | 36.2 | 905 | 141 | 2,820 | | | 800 |
| RpN4 | May 2016 | 36.6 | 915 | 165 | 3,300 | | | 800 |
| RpN5 | June 2016 | 54.8 | 1370 | 197 | 3,940 | | | 800 |
| RpN6 | July 2016 | 26 | 650 | 103 | 2,060 | | | 800 |
| RpN7 | July 2016 | 22.2 | 555 | 172 | 3,440 | | | 800 |
| RpN8 | July 2016 | 26.4 | 660 | 170 | 3,400 | | | 800 |
| RpN9 | July 2016 | 56 | 1400 | 152 | 3,040 | | | 800 |
| RpN10 | July 2016 | 20 | 500 | 128 | 2,560 | | | 800 |
| RpN11 | Oct 2016 | 10.2 | 255 | 136 | 2,720 | | | 800 |
| RpN12 | Oct 2016 | 9.8 | 245 | 177 | 3,540 | | | 800 |
| RpN13 | Oct 2016 | 27.7 | 692.5 | 156 | 3,120 | | | 800 |
| RpN14 | Oct 2016 | 19.9 | 497.5 | 113 | 2,260 | | | 800 |
| RpN15 | Oct 2016 | 7.44 | 186 | 102 | 2,040 | | | 800 |
| RpN16 | July 2013 | 0.492 | 12.3 | 8.7 | 208.8 | 42 | 840 | 630 |
| RpN17 | July 2013 | 7.92 | 198 | 56.6 | 1358.4 | | | 800 |
| RpN18 | July 2013 | 0.788 | 19.7 | 38.4 | 1228.8 | | | 768 |
| RpN19 | July 2013 | 6.28 | 157 | 46.4 | 1484.8 | | | 696 |
| RpN20 | July 2013 | 3.69 | 92.25 | 41.8 | 1003.2 | | | 627 |
| RpN21 | Oct 2013 | 0.732 | 18.3 | 30.6 | 979.2 | | | 612 |
| RpN22 | Oct 2013 | 2 | 50 | 32.4 | 1036.8 | | | 648 |
| RpN23 | Oct 2013 | 1.98 | 49.5 | 27.8 | 667.2 | | | 556 |
| RpN24 | Oct 2013 | 6.56 | 164 | 85 | 2040 | | | 800 |
| RpN25 | Oct 2013 | 0.836 | 20.9 | 22.4 | 537.6 | | | 537.6 |
| RpN26 | July 2010 | 1.9 | 47.5 | 38.6 | 926.4 | | | 772 |

| | | | | | | | | |
|---|---|---|---|---|---|---|---|---|
| RpN27 | July 2010 | 7.36 | 184 | 121 | 2904 | | | 800 |
| RpN28 | July 2010 | 5.48 | 137 | 110 | 2640 | | | 800 |
| RpN29 | July 2010 | 8.56 | 214 | 81 | 1944 | | | 800 |
| RpN30 | July 2010 | 4.64 | 116 | 43.2 | 1036.8 | | | 648 |
| RpN31 | Oct 2010 | 6.48 | 162 | 109 | 2616 | | | 800 |
| RpN32 | Oct 2010 | 1.3 | 32.5 | 17.9 | 572.8 | | | 447.5 |
| RpN33 | Oct 2010 | 11.1 | 277.5 | 120 | 2880 | | | 800 |
| RpN34 | Oct 2010 | 12.5 | 312.5 | 81.2 | 1948.8 | | | 800 |
| RpN35 | Oct 2010 | 7.88 | 197 | 89 | 2136 | | | 800 |
| RpN36 | July 2004 | 12.3 | 369 | 87.8 | 2107.2 | | | 1930 |
| RpN37 | July 2004 | 1.75 | 52.5 | 34.1 | 750.2 | | | 750.2 |
| RpN38 | July 2004 | 14.2 | 426 | | | | | 400 |
| RpN39 | July 2004 | 13.1 | 393 | 63.2 | 1516.8 | | | 1390 |
| RpN40 | July 2004 | 6.47 | 194.1 | 11.5 | 253 | > 60 | 1080 | 1080 |
| RpN50 | July 2010 | 22 | 660 | | | | | 616 |
| RpN51 | July 2007 | 17.8 | 534 | | | | | 498.4 |
| RpN52 | July 2007 | 10.5 | 315 | 22.7 | 544.8 | | | 500 |
| RpN53 | July 2007 | 16.4 | 492 | | | | | 460 |
| RpN54 | July 2007 | 44.4 | 1332 | | | | | 1243 |
| RpN55 | July 2007 | 9.43 | 282.9 | 22.6 | 542.4 | | | 497 |
| RpN56 | July 2013 | 0.696 | 20.88 | 7.43 | 178.32 | > 60 | 1080 | 1080 |
| RpN57 | July 2013 | 1.21 | 36.3 | 23.1 | 554.4 | | | 508 |
| RpN58 | July 2013 | 33.2 | 996 | | | | | 930 |
| RpN59 | July 2013 | 23.1 | 693 | | | | | 647 |
| RpN60 | July 2013 | 12.1 | 363 | | | | | 339 |
| RpN61 | July 2004 | 13.6 | 408 | | | | | 381 |
| RpN62 | July 2004 | 5.55 | 166.5 | 11.1 | 244.2 | > 60 | 1080 | 1080 |
| RpN64 | July 2004 | 6.66 | 199.8 | 42.6 | 766.8 | | | 937 |
| RpN66 | Aug 2007 | 7.09 | 212.7 | 58.7 | 1408.8 | | | 1291 |
| RpN68 | July 2007 | 23.8 | 714 | 23.8 | 714 | | | 666 |

| | | | | | | | | |
|---|---|---|---|---|---|---|---|---|
| RpN69 | July 2007 | 19.2 | 576 | 19.2 | 576 | | | 537.6 |
| RpN70 | Aug 2007 | 17.6 | 528 | 17.6 | 528 | | | 493 |
| RpN71 | July 2010 | 2.89 | 86.7 | 38.6 | 926.4 | | | 849 |
| RpN72 | July 2010 | 9.54 | 286.2 | > 60 | 1320 | | | 1320 |
| RpN73 | July 2010 | 7.55 | 226.5 | > 60 | 1320 | | | 1320 |
| RpN74 | July 2010 | 3.74 | 112.2 | 47.1 | 1130.4 | | | 1036 |
| RpN75 | July 2010 | 16.2 | 486 | | | | | 453.6 |
| RpN76 | July 2013 | 6.6 | 198 | 40.5 | 972 | | | 891 |
| RpN77 | July 2013 | 1.81 | 54.3 | 10 | 240 | > 60 | 1080 | 1080 |
| RpN78 | July 2013 | 12.2 | 366 | | | | | 341.6 |
| RpN80 | July 2013 | 2.62 | 78.6 | 18.8 | 451.2 | | | 413.6 |
| RpP1 | May 2016 | 7.08 | 177 | 57.6 | 1,152 | | | 800 |
| RpP2 | May 2016 | 6.96 | 174 | 40.8 | 816 | | | 612 |
| RpP3 | May 2016 | 25.1 | 627.5 | 141 | 2,820 | | | 800 |
| RpP4 | May 2016 | 3.43 | 85.75 | 2.32 | 46 | 184 | 3680 | 800 |
| RpP5 | May 2016 | 24.5 | 612.5 | 92.2 | 1,844 | | | 800 |
| RpP6 | July 2016 | 3.19 | 79.75 | 4.64 | 93 | 324 | 6480 | 800 |
| RpP7 | July 2016 | 7.04 | 176 | 4.18 | 84 | 244 | 4880 | 800 |
| RpP8 | July 2016 | 14.8 | 370 | 58.4 | 1,168 | | | 800 |
| RpP10 | July 2016 | 7.36 | 184 | 63.2 | 1,264 | | | 800 |
| RpP11 | Oct 2016 | 11.9 | 297.5 | 157 | 3,140 | | | 800 |
| RpP12 | Oct 2016 | 21.5 | 537.5 | 151 | 3,020 | | | 800 |
| RpP13 | Oct 2016 | 8.16 | 204 | 79.4 | 1,588 | | | 800 |
| RpP14 | Oct 2016 | 11.6 | 290 | 136 | 2,720 | | | 800 |
| RpP15 | Oct 2016 | 12.9 | 322.5 | 94.6 | 1,892 | | | 800 |
| RpP16 | July 2016 | 8.64 | 216 | 112 | 2,688 | | | 800 |
| RpP17 | July 2013 | 0.992 | 24.8 | 46.8 | 1497.6 | | | 702 |
| RpP18 | July 2013 | <0.01 | n.a. | 5.54 | 177.28 | 32.4 | 648 | 486 |
| RpP20 | July 2013 | 0.828 | 20.7 | 33.4 | 1068.8 | | | 668 |
| RpP21 | July 2013 | 1.49 | 37.25 | 54.8 | 1753.6 | | | 657.6 |

| | | | | | | | | |
|---|---|---|---|---|---|---|---|---|
| RpP22 | Oct 2013 | 3.1 | 77.5 | 50.6 | 1619.2 | | | 759 |
| RpP23 | Oct 2013 | <0.01 | n.a. | 22.2 | 710.4 | | | 555 |
| RpP24 | Oct 2013 | 0.892 | 22.3 | 32.4 | 1036.8 | | | 648 |
| RpP25 | Oct 2013 | 0.512 | 12.8 | 21.6 | 691.2 | | | 540 |
| RpP27 | July 2010 | 3.52 | 88 | 53.4 | 1708.8 | | | 800 |
| RpP28 | July 2010 | 0.58 | 14.5 | 19.6 | 627.2 | | | 490 |
| RpP29 | July 2010 | 4.88 | 122 | 54.8 | 1753.6 | | | 800 |
| RpP30 | July 2010 | <0.01 | n.a. | 17.6 | 563.2 | | | 563.2 |
| RpP31 | July 2010 | <0.01 | n.a. | 19.8 | 633.6 | | | 495 |
| RpP32 | Oct 2010 | 3.41 | 85.25 | 41.8 | 1337.6 | | | 627 |
| RpP33 | Oct 2010 | <0.01 | n.a. | 14.6 | 467.2 | 31.8 | 636 | 636 |
| RpP34 | Oct 2010 | 1.09 | 27.25 | 40.8 | 1305.6 | | | 612 |
| RpP35 | Oct 2010 | 0.952 | 23.8 | 19.6 | 627.2 | | | 490 |
| RpP36 | Oct 2010 | 1.45 | 36.25 | 22.6 | 723.2 | | | 565 |
| RpP37 | July 2013 | 1.04 | 26 | 7.48 | 239.36 | 35.8 | 859.2 | 537 |
| RpP40 | Oct 2013 | 3.22 | 80.5 | 41.4 | 1324.8 | | | 621 |
| RpSX1 | Apr 2016 | 4.54 | 90.8 | 123 | 2,460 | | | 800 |
| RpSX3 | May 2016 | 7.92 | 158.4 | 143 | 2,860 | | | 800 |
| RpSX9 | July 2016 | 1.04 | 20.8 | 5.98 | 120 | 312 | 6240 | 800 |
| RpSX11 | Oct 2016 | 2.95 | 59 | 39.6 | 792 | 498 | 9960 | 800 |
| RpSX13 | Oct 2016 | 4.03 | 80.6 | 46.8 | 936 | 470 | 9400 | 800 |
| RpSX16 | July 2013 | <0.01 | n.a. | 10.8 | 346 | | | 345.6 |
| RpSX17 | July 2013 | 0.984 | 19.68 | 34.2 | 1,094 | | | 684 |
| RpSX24 | Oct 2013 | 0.608 | 30.4 | 27 | 864 | | | 675 |
| RpSX27 | July 2010 | 0.548 | 27.4 | 39.8 | 1,274 | | | 796 |
| RpSX31 | Oct 2010 | 0.852 | 21.3 | 64 | 2,048 | | | 800 |
| RpSX32 | Oct 2010 | 0.684 | 17.1 | 47.4 | 1,517 | | | 711 |
| RpSX34 | Oct 2010 | 2.02 | 50.5 | 40 | 1,280 | | | 800 |
| RpSX35 | Oct 2010 | 0.788 | 19.7 | 23.6 | 755 | | | 590 |
| RpSX37 | May 2016 | 6.72 | 168 | 124 | 2,976 | | | 800 |

| | | | | | | | | |
|---|---|---|---|---|---|---|---|---|
| RpSX38 | May 2016 | 15.2 | 380 | 135 | 3,240 | | | 800 |
| RpSX39 | May 2016 | 6.32 | 158 | 125 | 3,000 | | | 800 |
| RpSX40 | July 2016 | 1.08 | 27 | 82 | 1,968 | | | 800 |
| RpSX41 | July 2016 | 5.2 | 130 | 117 | 2,808 | | | 800 |
| RpSX42 | July 2016 | 2.93 | 73.25 | 92.6 | 2,222 | | | 800 |
| RpSX43 | July 2016 | 1.3 | 32.5 | 73.2 | 1,757 | | | 800 |
| RpSX44 | Oct 2016 | 3.22 | 80.5 | 90 | 2,160 | | | 800 |
| RpSX45 | Oct 2016 | 6.96 | 174 | 126 | 3,024 | | | 800 |
| RpSX46 | Oct 2016 | 8.08 | 202 | 117 | 2,808 | | | 800 |
| RpSX49 | July 2013 | <0.01 | n.a. | 4.4 | 140.8 | 338 | 6760 | 800 |
| RpSX50 | July 2013 | <0.01 | n.a. | 9.06 | 289.92 | 140 | 2800 | 800 |
| RpSX51 | July 2013 | <0.01 | n.a. | 10.7 | 342.4 | 414 | 8280 | 800 |
| RpSX52 | Oct 2013 | 5.64 | 141 | 55.4 | 1772.8 | | | 831 |
| RpSX54 | Oct 2013 | 1.45 | 36.25 | 49.8 | 1593.6 | | | 747 |
| RpSX55 | Oct 2013 | 3 | 75 | 56.6 | 1811.2 | | | 800 |
| RpSX56 | Oct 2013 | 0.484 | 12.1 | 26 | 832 | | | 520 |
| RpSX58 | July 2010 | <0.01 | n.a. | 12.4 | 396.8 | 222 | 4440 | 800 |
| RpSX59 | July 2010 | 0.556 | 13.9 | 2.36 | 75.52 | 45 | 900 | 675 |
| RpSX60 | July 2010 | <0.01 | n.a. | 15.8 | 505.6 | 88.6 | 1772 | 800 |
| RpSX64 | Oct 2010 | 1.86 | 46.5 | 27.2 | 870.4 | | | 680 |
| RpSX66 | July 2010 | <0.01 | n.a. | 3.9 | 124.8 | 54.2 | 1300.8 | 813 |
| RpSX68 | July 2004 | 0.794 | 23.82 | >60 | 1320 | | | 1320 |
| RpSX69 | July 2004 | 5.75 | 172.5 | >60 | 1320 | | | 1320 |
| RpSX70 | July 2004 | 6.56 | 196.8 | 50 | 1200 | | | 1100 |
| RpSX71 | July 2004 | 2.76 | 82.8 | 34.8 | 835.2 | | | 765.6 |
| RpSX72 | July 2013 | 1.88 | 56.4 | 33.2 | 796.8 | | | 1144 |
| RpSX73 | July 2013 | 1.66 | 49.8 | 24.1 | 578.4 | | | 530.2 |
| RpSX74 | July 2013 | 0.722 | 21.66 | >60 | 1320 | | | 1276 |
| RpSX75 | July 2013 | 1.67 | 50.1 | 45.7 | 1096.8 | | | 532.4 |
| RpSX76 | July 2013 | 0.595 | 17.85 | 44.1 | 1058.4 | | | 1020 |

| | | | | | | |
|---|---|---|---|---|---|---|
| RpSX77 | July 2007 | 1.76 | 52.8 | 75.4 | 1809.6 | 1658.8 |
| RpSX78 | July 2007 | 0.62 | 18.6 | 53.2 | 1276.8 | 1170.4 |
| RpSX79 | July 2007 | 0.973 | 29.19 | 57 | 1368 | 1254 |
| RpSX80 | July 2007 | 0.697 | 20.91 | 55 | 1320 | 1210 |
| RpSX81 | July 2007 | 1.33 | 39.9 | >60 | 1320 | 1320 |
| RpSX82 | July 2010 | 0.303 | 9.09 | 34.3 | 823.2 | 754.6 |
| RpSX83 | July 2010 | 0.446 | 13.38 | 59 | 1416 | 1298 |
| RpSX84 | July 2010 | 0.455 | 13.65 | >60 | 1320 | 1320 |
| RpSX85 | July 2010 | 0.958 | 28.74 | 48.7 | 1168.8 | 1071.4 |
| RpSX86 | July 2010 | 0.721 | 21.63 | 43.2 | 1036.8 | 950.4 |
| RpSX87 | July 2004 | 6.01 | 180.3 | 49.7 | 1192.8 | 1093.4 |
| RpSX88 | July 2004 | 0.58 | 17.4 | 26.9 | 646 | 591.8 |
| RpSX89 | July 2004 | 1.83 | 54.9 | >60 | 1320 | 1320 |
| RpSX90 | July 2004 | 6.09 | 182.7 | 50 | 1200 | 1100 |
| RpSX91 | July 2004 | 5.33 | 159.9 | >60 | 1320 | 1320 |
| RpSX92 | July 2007 | 1.27 | 38.1 | 26.7 | 640.8 | 587.4 |
| RpSX93 | July 2007 | 1.12 | 33.6 | 25.4 | 609.6 | 558.8 |
| RpSX94 | July 2007 | 1.33 | 39.9 | >60 | 1320 | 1320 |
| RpSX95 | July 2007 | 1.25 | 37.5 | 55 | 1320 | 1210 |
| RpSX96 | July 2007 | 1.03 | 30.9 | 32.3 | 775.2 | 710.6 |
| RpSX97 | July 2010 | 0.692 | 20.76 | 31.8 | 763.2 | 699.6 |
| RpSX98 | July 2010 | 0.817 | 24.51 | 58 | 1392 | 1276 |
| RpSX99 | July 2010 | 0.674 | 20.22 | 39 | 936 | 858 |
| RpSX100 | July 2010 | 0.516 | 15.48 | 19.8 | 475.2 | 435.6 |
| RpSX101 | July 2010 | 0.695 | 20.85 | 15.3 | 367.2 | 336.6 |
| RpSX103 | July 2013 | 0.37 | 11.1 | >60 | 1320 | 1320 |
| RpSX104 | July 2013 | 0.419 | 12.57 | 22 | 528 | 488.4 |
| RpSX105 | July 2013 | 0.624 | 18.72 | 13.9 | 333.6 | 320 |
| RpSX106 | July 2013 | 0.419 | 12.57 | >60 | 1320 | 794.2 |
| RpW1 | Apr 2016 | 16.1 | 402.5 | 220 | 4,400 | 800 |

| | | | | | | |
|---|---|---|---|---|---|---|
| RpW2 | May 2016 | 10.2 | 255 | 131 | 2,620 | 800 |
| RpW3 | May 2016 | 7.68 | 192 | 82.6 | 1,652 | 800 |
| RpW4 | May 2016 | 9.92 | 248 | 168 | 3,360 | 800 |
| RpW5 | May 2016 | 21.8 | 545 | 212 | 4,240 | 800 |
| RpW6 | July 2016 | 5 | 125 | 119 | 2,380 | 800 |
| RpW7 | July 2016 | 8.4 | 210 | 177 | 3,540 | 800 |
| RpW8 | July 2016 | 6.1 | 152.5 | 97.2 | 1,944 | 800 |
| RpW9 | July 2016 | 16.1 | 402.5 | 208 | 4,160 | 800 |
| RpW10 | July 2016 | 18.4 | 460 | 206 | 4,120 | 800 |
| RpW11 | Oct 2016 | 18.4 | 460 | 248 | 4,960 | 800 |
| RpW12 | Oct 2016 | 16.7 | 417.5 | 258 | 5,160 | 800 |
| RpW13 | Oct 2016 | 15.9 | 397.5 | 230 | 4,600 | 800 |
| RpW14 | Oct 2016 | 12.6 | 315 | 206 | 4,120 | 800 |
| RpW15 | Oct 2016 | 8.68 | 217 | 188 | 3,760 | 800 |
| RpWr3 | July 2016 | 17.9 | 537 | | | 501.2 |
| RpWr9 | July 2016 | 14.1 | 423 | | | 395 |
| RpWr12 | July 2016 | 25.7 | 771 | | | 719.6 |
| RpWr13 | July 2016 | 18.4 | 552 | | | 515.2 |
| RpWr14 | July 2016 | 20.6 | 618 | | | 576.8 |
| RpWr15 | July 2016 | 28.6 | 858 | | | 800.8 |
| RpWr16 | Aug 2016 | 17.4 | 522 | | | 487.2 |
| RpWr19 | Aug 2016 | 18.7 | 561 | | | 523.6 |
| RpWr22 | July 2016 | 16.7 | 501 | | | 467.6 |
| RpWr23 | July 2016 | 21.3 | 639 | | | 596.4 |
| RpY6 | July 2016 | 21.7 | 651 | | | 607.6 |
| RpY8 | July 2016 | 16.6 | 498 | | | 464.8 |
| RpY9 | July 2016 | 16.1 | 483 | | | 450.8 |
| RpY11 | July 2016 | 18.2 | 546 | | | 509.6 |
| RpY14 | Aug 2016 | 19.3 | 579 | | | 540.4 |
| RpY16 | Aug 2016 | 19.6 | 588 | | | 548.8 |

| | | | | |
|---|---|---|---|---|
| RpY21 | Aug 2016 | 35.3 | 1059 | 988.4 |
| RpY22 | Aug 2016 | 17.6 | 528 | 492.8 |
| RpY26 | Aug 2016 | 16.6 | 498 | 464.8 |
| RpY27 | Aug 2016 | 33.8 | 1014 | 946.4 |

**Table S3.** Filtering schemes for the different data sets used in the population structure analyses: A) all samples included; B) samples from Newcastle only; C) samples from Starcross only. The order of rows indicates the sequential filters applied to the data. minDP – include only genotypes with depth greater or equal to the value; minQ – include sites with quality above the value; mac – include sites with minor allele count greater or equal to the value; geno – retain sites that have been successfully genotyped in the given proportion of individuals (max-missing filtering option in vcftools); imiss – retain individuals with a proportion of missing data smaller than the value. FS highlighted in green show the datasets used in the analyses.

**A)**

| FILTER | FS1 | FS2 | FS3 | FS4 | FS5 | FS6 | FS7 |
|---|---|---|---|---|---|---|---|
| **MISSING DATA** | | | | | | geno > 50% | geno > 50%<br>remove-indels |
| **LOW-CONFIDENCE SNP CALL** | | | minDP > 5<br>minQ > 20<br>mac > 3 | minDP > 5<br>minQ > 20<br>mac > 3 | minDP > 5<br>minQ > 20<br>mac > 3 | mac > 3<br>minQ > 30<br>minDP > 3 | mac >3<br>minQ > 20<br>minDP > 3 |
| **MISSING DATA** | geno > 90%<br>remove-indels<br>imiss < 25% / < 50%<br>Not done imiss for all samples as it is not worthy | imiss < 78%<br>geno > 90% | geno > 90% | imiss < 90%<br>geno > 90% | geno > 15%<br>imiss < 95%<br>geno > 60%<br>imiss < 80%<br>geno > 70%<br>imiss < 75%<br>geno > 80% | imiss < 95%<br>geno > 75% | imiss < 60%<br>geno > 90% |
| **LOW-CONFIDENCE SNP CALL** | | minDP > 5<br>minQ > 20<br>mac > 3 | | | | | minDP > 5 |
| **MISSING DATA** | | | | | | | geno > 95% |
| **INFO FILTERS** | Remove-indels | Remove-indels | | | | remove-indels | thin 2000 |
| **SNPS** | 2 | 63,187 (2,287,871) | 0 | 1704 | 3244 (2,287,871) | 1526 | 4802 |
| **INDIVIDUALS** | Not done | 84 (175) | 175 | 60 (175) | 67 (175) | 91 | 86 |

**B)**

| FILTER | FS1 | FS2 | FS3 | FS4 |
|---|---|---|---|---|
| **MISSING DATA** | geno > 50%<br>remove-indels | remove-indels<br>geno > 50% | geno > 50%<br>imiss < 99% | |

| | | | | |
|---|---|---|---|---|
| | | | geno > 75% | |
| **LOW-CONFIDENCE SNP CALL** | | min-meanDP > 5<br>mac > 3<br>minQ > 20 | minDP > 3<br>mac > 3<br>minQ > 20 | minDP > 3<br>mac > 3<br>minQ > 20 |
| **MISSING DATA** | imiss < 75%<br>geno > 90%<br><br>thin 2Kb | imiss < 95%<br>geno > 80%<br><br>thin 2kb | geno > 50%<br>thin 2kb<br>remove-indels<br>imiss < 90% | thin 2kb<br>geno > 30%<br>imiss < 99%<br>remove-indels<br>imiss < 95<br>geno > 60%<br>geno > 70% |
| **LOW-CONFIDENCE SNP CALL** | min-meanDP > 5<br>mac > 3<br>minQ > 20 | | | |
| **MISSING DATA** | | | | |
| **INFO FILTERS** | | | | |
| **SNPS** | 1746 (< 10% missing per locus) | 3186 (< 20% missing per locus) | 5277 (< 50% missing per locus) | 359 (< 30% missing per locus) |
| **INDIVIDUALS** | 28 (< 50% missing per indv) | 36 (0-90% missing per indv) | 29 (0-85% missing per indv) | 33 (0 – 91% missing per indv) |

**C)**

| **FILTER** | **FS1** | **FS2** | **FS3** | **FS4** |
|---|---|---|---|---|
| **MISSING DATA** | geno > 50%<br>remove-indels | remove-indels<br>geno > 25% | geno > 25%<br>imiss < 99%<br>geno > 50% | |
| **LOW-CONFIDENCE SNP CALL** | | min-meanDP > 5<br>mac > 3<br>minQ > 20 | minDP > 3<br>mac > 3<br>minQ > 20 | minDP > 3<br>mac > 3<br>minQ > 20 |
| **MISSING DATA** | imiss < 95%<br>thin 2kb | imiss < 95%<br>geno > 70%<br>think 2K | geno > 60%<br>thin 2kb<br>remove-indels<br>imiss < 90% | thin 2kb<br>geno > 20%<br>imiss < 99%<br>remove-indels |

| | | | | |
|---|---|---|---|---|
| | | | | geno > 60%<br>imiss < 95% |
| **LOW-CONFIDENCE SNP CALL** | minDP > 3<br>mac > 3<br>minQ > 20 | | | |
| **MISSING DATA** | | | | |
| **INFO FILTERS** | | | | |
| **SNPS** | 89 (< 90% missing per locus) | 907 (< 30% missing per locus) | 1638 (< 40% missing data per locus) | 359 (< 50% missing per locus) |
| **INDIVIDUALS** | 46 (0-99% missing per indv) | 31 (0-89% missing data per indv) | 18 (< 75% missing data per indv) | 27 (0-92% missing per indv) |

**Table S4.** Sequencing results for the aphids used in GBS showing the number of reads that passed the QC, the number and percentage of reads aligned to the *R. padi* genome.

| Sample | Trap date | N reads | Mapped reads | Percentage mapped |
|---|---|---|---|---|
| RpN1 | May 2016 | 4995472 | 4954101 | 99.17% |
| RpN10 | July 2016 | 5200744 | 4826984 | 92.81% |
| RpN11 | October 2016 | 5167156 | 5130989 | 99.30% |
| RpN12 | October 2016 | 5714318 | 5675868 | 99.33% |
| RpN13 | October 2016 | 8999479 | 8936557 | 99.30% |
| RpN14 | October 2016 | 8738807 | 8672698 | 99.24% |
| RpN15 | October 2016 | 9016847 | 8580189 | 95.16% |
| RpN16 | July 2013 | 1961831 | 87707 | 4.47% |
| RpN2 | May 2016 | 4819781 | 4786969 | 99.32% |
| RpN23 | October 2013 | 339022 | 252469 | 74.47% |
| RpN26 | July 2010 | 276 | 0 | 0.00% |
| RpN27 | July 2010 | 6925877 | 6713675 | 96.94% |
| RpN28 | July 2010 | 8563521 | 8464873 | 98.85% |
| RpN29 | July 2010 | 6146045 | 6075966 | 98.86% |
| RpN3 | May 2016 | 5244950 | 5066806 | 96.60% |
| RpN30 | July 2010 | 2835505 | 2449152 | 86.37% |
| RpN31 | October 2010 | 10039372 | 9896932 | 98.58% |
| RpN33 | October 2010 | 9595798 | 9258270 | 96.48% |
| RpN34 | October 2010 | 9714165 | 9614719 | 98.98% |
| RpN36 | July 2004 | 5287189 | 5118021 | 96.80% |
| RpN37 | July 2004 | 1611708 | 188437 | 11.69% |
| RpN38 | July 2004 | 3192066 | 3154267 | 98.82% |
| RpN39 | July 2004 | 3301 | 504 | 15.27% |
| RpN4 | May 2016 | 5079105 | 5028279 | 99.00% |
| RpN40 | July 2004 | 101592 | 65727 | 64.70% |
| RpN5 | June 2016 | 4123268 | 1507762 | 36.57% |
| RpN50 | July 2010 | 1864 | 665 | 35.68% |
| RpN51 | July 2007 | 2130607 | 2092445 | 98.21% |
| RpN53 | July 2007 | 1377 | 459 | 33.33% |
| RpN54 | July 2007 | 4018278 | 3988600 | 99.26% |
| RpN55 | July 2007 | 2173309 | 1969894 | 90.64% |
| RpN56 | July 2013 | 3186214 | 2645666 | 83.03% |
| RpN57 | July 2013 | 4055049 | 3568421 | 88.00% |
| RpN58 | July 2013 | 500 | 184 | 36.80% |
| RpN59 | July 2013 | 652144 | 631898 | 96.90% |
| RpN6 | July 2016 | 8208385 | 8131345 | 99.06% |
| RpN60 | July 2013 | 1209 | 729 | 60.30% |
| RpN61 | July 2004 | 2463628 | 2440920 | 99.08% |
| RpN62 | July 2004 | 1138684 | 45692 | 4.01% |
| RpN64 | July 2004 | 3113092 | 2249116 | 72.25% |
| RpN68 | July 2007 | 1900239 | 1882661 | 99.07% |
| RpN69 | July 2007 | 1753805 | 1740003 | 99.21% |

| | | | | |
|---|---|---|---|---|
| RpN7 | July 2016 | 7854931 | 6955765 | 88.55% |
| RpN70 | August 2007 | 1591 | 601 | 37.77% |
| RpN71 | July 2010 | 5065311 | 228746 | 4.52% |
| RpN72 | July 2010 | 3362448 | 2508245 | 74.60% |
| RpN73 | July 2010 | 4570586 | 3943462 | 86.28% |
| RpN74 | July 2010 | 4064388 | 2691465 | 66.22% |
| RpN75 | July 2010 | 1300 | 1228 | 94.46% |
| RpN76 | July 2013 | 3733310 | 2027461 | 54.31% |
| RpN77 | July 2013 | 1224125 | 14736 | 1.20% |
| RpN78 | July 2013 | 1206 | 666 | 55.22% |
| RpN8 | July 2016 | 6336591 | 6207972 | 97.97% |
| RpN80 | July 2013 | 3910976 | 62157 | 1.59% |
| RpN9 | July 2016 | 7856312 | 7751657 | 98.67% |
| RpP1 | May 2016 | 5702474 | 5635118 | 98.82% |
| RpP10 | July 2016 | 728 | 3 | 0.41% |
| RpP11 | October 2016 | 6028232 | 5956261 | 98.81% |
| RpP12 | October 2016 | 220 | 1 | 0.45% |
| RpP13 | October 2016 | 4724102 | 4471533 | 94.65% |
| RpP14 | October 2016 | 5222045 | 5155751 | 98.73% |
| RpP15 | October 2016 | 4580228 | 4367378 | 95.35% |
| RpP16 | July 2016 | 7669657 | 7605084 | 99.16% |
| RpP2 | May 2016 | 4120232 | 4081665 | 99.06% |
| RpP22 | October 2013 | 3948582 | 3500291 | 88.65% |
| RpP27 | July 2010 | 462 | 1 | 0.22% |
| RpP28 | July 2010 | 2484910 | 645630 | 25.98% |
| RpP29 | July 2010 | 3349768 | 3097709 | 92.48% |
| RpP3 | May 2016 | 5774855 | 5736933 | 99.34% |
| RpP30 | July 2010 | 12 | 0 | 0.00% |
| RpP31 | July 2010 | 30 | 0 | 0.00% |
| RpP4 | May 2016 | 6570900 | 6217932 | 94.63% |
| RpP40 | October 2013 | 5735515 | 5231599 | 91.21% |
| RpP5 | May 2016 | 3588698 | 3265486 | 90.99% |
| RpP6 | July 2016 | 4951705 | 2230109 | 45.04% |
| RpP7 | July 2016 | 7097326 | 2766864 | 38.98% |
| RpP8 | July 2016 | 5075651 | 4739580 | 93.38% |
| RpSX1 | April 2016 | 36 | 1 | 2.78% |
| RpSX100 | July 2010 | 2681980 | 12496 | 0.47% |
| RpSX101 | July 2010 | 1314275 | 54636 | 4.16% |
| RpSX103 | July 2013 | 1513767 | 45294 | 2.99% |
| RpSX104 | July 2013 | 395897 | 28184 | 7.12% |
| RpSX105 | July 2013 | 3080947 | 48005 | 1.56% |
| RpSX106 | July 2013 | 2581532 | 8291 | 0.32% |
| RpSX11 | October 2016 | 2357899 | 2323869 | 98.56% |
| RpSX13 | October 2016 | 5910183 | 5766040 | 97.56% |
| RpSX24 | October 2013 | 5205780 | 89798 | 1.72% |
| RpSX27 | July 2010 | 2317087 | 1251627 | 54.02% |

| | | | | |
|---|---|---|---|---|
| RpSX3 | May 2016 | 5030871 | 4974208 | 98.87% |
| RpSX37 | May 2016 | 6389272 | 5198434 | 81.36% |
| RpSX38 | May 2016 | 7059250 | 7010900 | 99.32% |
| RpSX39 | May 2016 | 3568984 | 3518168 | 98.58% |
| RpSX40 | July 2016 | 6079061 | 4970569 | 81.77% |
| RpSX41 | July 2016 | 7696800 | 7433991 | 96.59% |
| RpSX42 | July 2016 | 9045117 | 8886832 | 98.25% |
| RpSX43 | July 2016 | 6952809 | 3405164 | 48.98% |
| RpSX44 | October 2016 | 6070685 | 5919963 | 97.52% |
| RpSX45 | October 2016 | 6235064 | 5914984 | 94.87% |
| RpSX46 | October 2016 | 5714428 | 5628214 | 98.49% |
| RpSX52 | October 2013 | 7703013 | 7259462 | 94.24% |
| RpSx54 | October 2013 | 6777513 | 6360915 | 93.85% |
| RpSX55 | October 2013 | 7199403 | 6699893 | 93.06% |
| RpSX56 | October 2013 | 5315374 | 3657532 | 68.81% |
| RpSX58 | July 2010 | 14 | 0 | 0.00% |
| RpSX59 | July 2010 | 507165 | 4961 | 0.98% |
| RpSX60 | July 2010 | 2876233 | 20172 | 0.70% |
| RpSX64 | October 2010 | 2513020 | 2023064 | 80.50% |
| RpSX66 | July 2010 | 6692363 | 577943 | 8.64% |
| RpSX68 | July 2004 | 1272256 | 73603 | 5.79% |
| RpSX69 | July 2004 | 2401744 | 1322919 | 55.08% |
| RpSX70 | July 2004 | 3314172 | 281289 | 8.49% |
| RpSX71 | July 2004 | 4365532 | 544744 | 12.48% |
| RpSX72 | July 2013 | 2412080 | 1132019 | 46.93% |
| RpSX74 | July 2013 | 667371 | 27545 | 4.13% |
| RpSX75 | July 2013 | 245196 | 62780 | 25.60% |
| RpSX76 | July 2013 | 739755 | 7057 | 0.95% |
| RpSX77 | July 2007 | 202720 | 16019 | 7.90% |
| RpSX78 | July 2007 | 532 | 16 | 3.01% |
| RpSX79 | July 2007 | 1085301 | 324270 | 29.88% |
| RpSX80 | July 2007 | 764577 | 288423 | 37.72% |
| RpSX81 | July 2007 | 770022 | 48394 | 6.28% |
| RpSX82 | July 2010 | 1573104 | 111450 | 7.08% |
| RpSX83 | July 2010 | 341770 | 35717 | 10.45% |
| RpSX84 | July 2010 | 1970780 | 33920 | 1.72% |
| RpSX85 | July 2010 | 1316713 | 600969 | 45.64% |
| RpSX86 | July 2010 | 4019654 | 3163892 | 78.71% |
| RpSX87 | July 2004 | 6052227 | 4697286 | 77.61% |
| RpSX88 | July 2004 | 6848215 | 4988476 | 72.84% |
| RpSX89 | July 2004 | 3297635 | 59740 | 1.81% |
| RpSX9 | July 2016 | 8279828 | 7767546 | 93.81% |
| RpSX90 | July 2004 | 2916365 | 1644707 | 56.40% |
| RpSX91 | July 2004 | 3215721 | 1395703 | 43.40% |
| RpSX92 | July 2007 | 11973 | 8607 | 71.89% |
| RpSX93 | July 2007 | 889 | 787 | 88.53% |

| | | | | |
|---|---|---|---|---|
| RpSX94 | July 2007 | 1879146 | 856579 | 45.58% |
| RpSX95 | July 2007 | 2014436 | 62310 | 3.09% |
| RpSX96 | July 2007 | 1121611 | 711733 | 63.46% |
| RpSX97 | July 2010 | 2404596 | 7163 | 0.30% |
| RpSX98 | July 2010 | 2810450 | 63968 | 2.28% |
| RpSX99 | July 2010 | 1982435 | 11898 | 0.60% |
| RpW1 | April 2016 | 6741040 | 6679197 | 99.08% |
| RpW10 | July 2016 | 6681470 | 6369210 | 95.33% |
| RpW11 | October 2016 | 5196967 | 5147924 | 99.06% |
| RpW12 | October 2016 | 5789812 | 5747978 | 99.28% |
| RpW13 | October 2016 | 6485111 | 6436316 | 99.25% |
| RpW14 | October 2016 | 6306791 | 6264912 | 99.34% |
| RpW15 | October 2016 | 5511667 | 5463819 | 99.13% |
| RpW2 | May 2016 | 5344539 | 4502008 | 84.24% |
| RpW3 | May 2016 | 2126278 | 1156035 | 54.37% |
| RpW4 | May 2016 | 6256548 | 6210680 | 99.27% |
| RpW5 | May 2016 | 5133083 | 5090163 | 99.16% |
| RpW6 | July 2016 | 7284018 | 6302234 | 86.52% |
| RpW7 | July 2016 | 7782377 | 6500144 | 83.52% |
| RpW8 | July 2016 | 7759623 | 4786033 | 61.68% |
| RpW9 | July 2016 | 6941326 | 6621067 | 95.39% |
| RpWr12 | July 2016 | 2025720 | 2008870 | 99.17% |
| RpWr13 | July 2016 | 1896207 | 1881431 | 99.22% |
| RpWr14 | July 2016 | 2433675 | 2414126 | 99.20% |
| RpWr15 | July 2016 | 1911181 | 1896378 | 99.23% |
| RpWr16 | August 2016 | 1811875 | 1797407 | 99.20% |
| RpWr19 | August 2016 | 1870903 | 1855521 | 99.18% |
| RpWr22 | July 2016 | 1513893 | 1496810 | 98.87% |
| RpWr23 | July 2016 | 2055206 | 2035664 | 99.05% |
| RpWr3 | July 2016 | 2220158 | 2117264 | 95.37% |
| RpWr9 | July 2016 | 1943112 | 1845462 | 94.97% |
| RpY11 | July 2016 | 1792378 | 1778307 | 99.21% |
| RpY14 | August 2016 | 2216720 | 2103338 | 94.89% |
| RpY16 | August 2016 | 1833230 | 1816076 | 99.06% |
| RpY21 | August 2016 | 5057783 | 4855700 | 96.00% |
| RpY22 | August 2016 | 1945851 | 1834913 | 94.30% |
| RpY26 | August 2016 | 2115759 | 2096472 | 99.09% |
| RpY27 | August 2016 | 5958699 | 5912133 | 99.22% |
| RpY6 | July 2016 | 3173082 | 3140742 | 98.98% |
| RpY8 | July 2016 | 2026918 | 2007246 | 99.03% |
| RpY9 | July 2016 | 1943929 | 1922756 | 98.91% |
| **Average** | | 3708552 | 2985915 | 65% |

**Table S5.** Pairwise genetic differentiation ($F_{ST}$) between samples from the North (A) and South (B) locations collected at different seasons in the year. N – Newcastle, P – Preston, Sx – Starcross, W – Wye. Column Ng shows the number of gene copies per sample group (number individuals x 2). Significant values are shown in italics.

**A)**

| | **Ng** | **N spring** | **N summer** | **N autumn** | **P spring** | **P summer** | **P autumn** |
|---|---|---|---|---|---|---|---|
| **N spring** | 8 | - | | | | | |
| **N summer** | 30 | *0.057* | - | | | | |
| **N autumn** | 16 | *0.067* | *0.039* | - | | | |
| **P spring** | 10 | 0.030 | 0.028 | *0.051* | - | | |
| **P summer** | 6 | 0.052 | 0.011 | *0.048* | 0.025 | - | |
| **P autumn** | 8 | *0.073* | *0.061* | *0.042* | *0.061* | *0.063* | - |

**B)**

| | **Ng** | **W spring** | **W summer** | **W autumn** | **Sx spring** | **Sx summer** | **Sx autumn** |
|---|---|---|---|---|---|---|---|
| **W spring** | 10 | - | | | | | |
| **W summer** | 10 | -0.038 | - | | | | |
| **W autumn** | 10 | *0.164* | *0.123* | - | | | |
| **Sx spring** | 8 | 0.032 | 0.003 | *0.235* | - | | |
| **Sx summer** | 6 | -0.023 | -0.050 | *0.082* | *0.079* | - | |
| **Sx autumn** | 10 | 0.005 | -0.013 | *0.102* | 0.040 | -0.020 | - |
